# Structural Determinants of Mucins in Influenza Virus Inhibition: The Synergistic Role of Sialylated Glycans and Molecular Size

**DOI:** 10.1101/2024.12.09.627521

**Authors:** Cosmin Butnarasu, Marc Safferthal, Jolly Thomas, Tatyana L. Povolotsky, Robyn Diehn, Kerstin Fentker, Philipp Mertins, Kevin Pagel, Daniel Lauster

## Abstract

Mucins are heavily glycosylated proteins that play a crucial role in protecting mucosal surfaces against pathogens, including influenza viruses. This study investigates the antiviral properties of bovine submaxillary mucins (BSM) as a model for oral mucins against the influenza virus (A/H3N2 subtype), focusing on glycan composition and mucin size. BSM was purified, and characterized by proteomic and glycomic analysis and its antiviral efficacy was assessed after selective removal of sialic acids, *N*-glycans, or all glycans via enzymatic and chemical treatments. We employed virus binding and inhibition assays, including microscale thermophoresis (MST) and hemagglutination inhibition (HAI), to characterize processed mucins for structure activity correlations. Removal of sialic acids reduced BSM’s antiviral activity by over 10-fold, while complete glycan removal abolished it entirely, highlighting sialylated *O*-glycans as critical for viral inhibition. *N*-glycan removal had minimal impact on antiviral efficacy. A size-dependent antiviral effect was observed: smaller mucin fragments (∼50 and 330 kDa), which retained comparable *O*-glycosylation patterns, showed significantly reduced inhibition and viral binding affinity compared to intact BSM. These findings underscore the importance of mucin size and sialylated *O*-glycans in antiviral defense mechanisms against influenza.

**Significance:** This study sheds light on the intrinsic antiviral properties of mucin from the bovine submaxillary gland, revealing the roles of glycans and mucin’s size in binding and inhibiting influenza virus. Our findings suggest a clear correlation between sialylated *O*-glycans and mucin size with the antiviral efficacy. Ultimately, we show that mucin-derived fragments retaining virus-binding capacity, with defined size and *O*-glycosylation patterns, can be isolated from mucin and could serve as versatile building blocks for designing next-generation antiviral biomaterials.

## 1. Introduction

Influenza A (IAV) infections continue to represent a major threat to global public health and economics. Seasonal influenza infects as many as 1 billion people, making it one of the most common respiratory diseases. The World Health Organization (WHO) estimates annually 3–5 million cases of severe illness and up to 650,000 deaths worldwide (1). Despite advances in vaccination strategies and antiviral therapies, the constant evolution of influenza viruses continues to pose challenges to effectively combat these pathogens (2).

One promising avenue for influenza prevention and treatment lies in understanding the intricate interplay between viral components and host factors during the initial steps of infection (3). Among these host factors, mucins, a family of heavily glycosylated proteins, represents a pivotal players in the defense against viral infections (4–6). As the main constituents of mucus, mucins provide protection by acting as both physical and chemical barriers to pathogens while maintaining the hydration of mucosal surfaces. The core protein is rich in serine and threonine, which can carry *O*-glycans and can make up to 80% of the mass of the protein. This extensive glycosylation gives mucins their characteristic bottlebrush-like structure, which is flexible and negatively charged overall (7).

In the initial stage of host cell infection, the viral hemagglutinin (HA) mediates the binding to sialic acid (SA) on the cell surface in a multivalent fashion, which leads to uptake of the virus particle into host cells (**Figure 1a**) (8–10). *N*-acetylneuraminic acid (Neu5Ac) and *N*-glycolylneuraminic acid (Neu5Gc) are the two major forms of SA relevant for influenza infections. While Neu5Ac is the predominant form in humans, Neu5Gc is found in other animals, such as cows and pigs (11). Most human-adapted influenza viruses, including H1N1 and H3N2 strains, preferentially bind to Neu5Ac, while animal-adapted virus strains can bind also Neu5Gc, which is not present in humans. An earlier study found that Neu5Gc acts as a decoy receptor for such IAV strains, but does not permit cell entry (12). Although the mucus mesh size of approximately 120 nm permits influenza virus penetration, the highly abundant SA on mucins restricts viral diffusion (13). Mucins, in fact, display SA as part of *N*- and *O*-glycans, and most probably they have evolved SA as a decoy for viruses to get them trapped in mucus, thereby preventing cell infection (14). However, the role of sialylated *N*- and *O*-glycans on the interaction with the influenza spike proteins HA and neuraminidase (NA) is still under debate, and most probably strain dependent (10).

**Figure 1.**
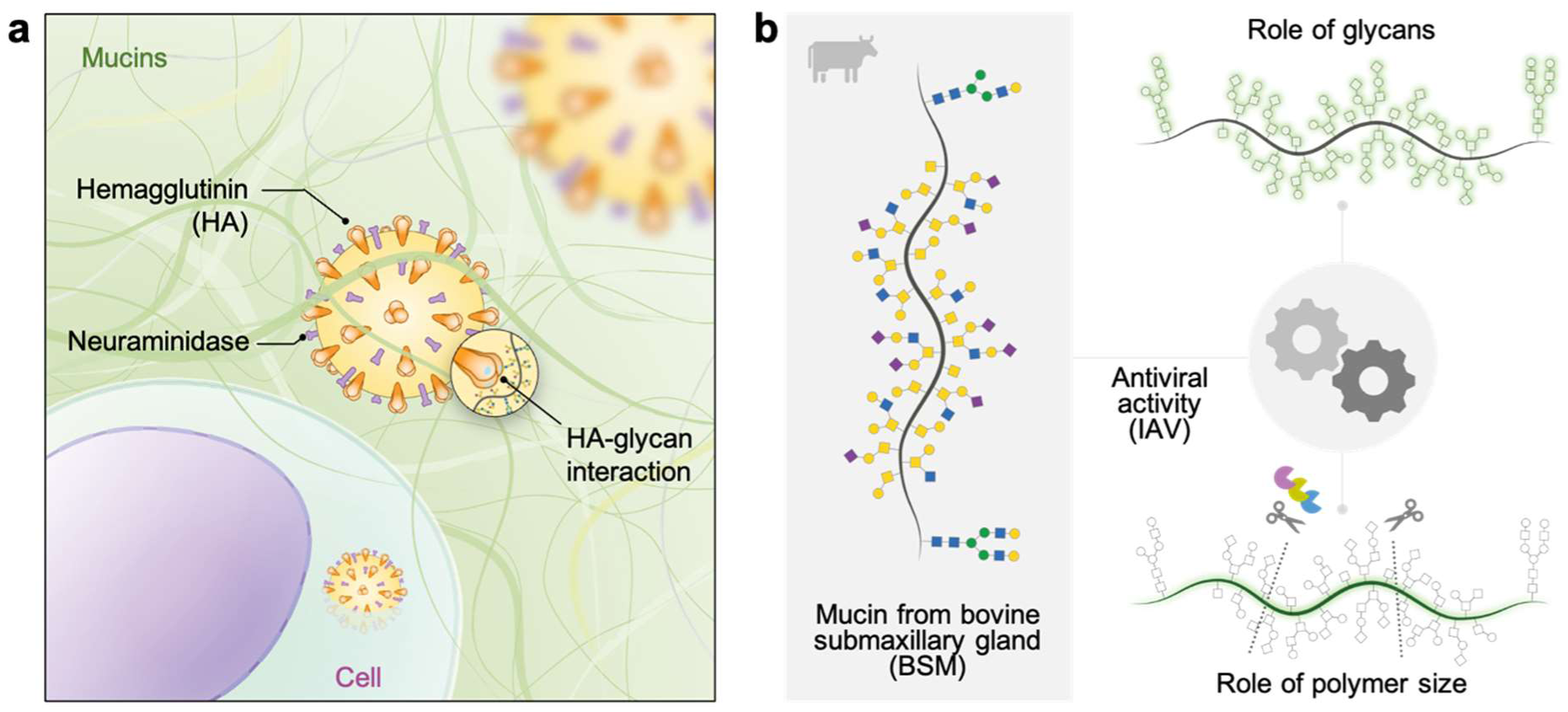
Mucins in the lungs play an important role as a barrier to viral infections such as influenza A virus (IAV). (**a**) The glycans on mucins act as decoy receptors for IAV hemagglutinin binding, reducing the probability of epithelial cell infections. (**b**) Schematic illustration of the approach used in this study to explore the effect of glycans and the size of the mucin from bovine submaxillary gland (BSM) on the binding to IAV.

It is clear that mucins are multifunctional macromolecules, which fulfill besides antiviral properties also many more (15). A defining characteristic of mucins that shapes their mechanical and functional properties is their remarkable size. Mucin monomers can extend up to 500 nm in length (16), forming an intricate, three-dimensional network that sterically shields viral particles while multivalently presenting SA residues. This structural organization is particularly effective against influenza viruses, as hemagglutinin (HA) binding requires an optimal spacing of ∼4.5 nm for intra-trimeric interactions and up to 14 nm for inter-trimeric binding (17, 18). In principle, the extended size and network formation of mucins allow them to meet these spatial criteria, enhancing their ability to bind and trap viral particles. This natural defense mechanism has inspired the design of synthetic biopolymers to inhibit viral infections mimicking the multivalent presentation of sialic acids on the host cell surface to block virus binding (19–22). Efforts to elucidate the impact of polymer structure on the inhibitory effect have shown, for instance, that linear polymers are superior to dendritic in inhibiting the influenza virus, highlighting the critical influence of polymer architecture (18). However, replicating the complex glycosylation patterns of natural mucins remains a significant challenge in synthetic chemistry (23) as well as in recombinant biology (24). Despite these advances, the size-dependent antiviral properties of natural mucins and their underlying biophysical mechanisms have yet to be systematically explored.

Here, we address the question of mucin’s antiviral activity by focusing on bovine submaxillary gland mucins (BSM). We selected BSM for its abundance of both Neu5Ac and Neu5Gc, with a predominance of Neu5Ac (25, 26). To investigate the role of glycan structures and size in antiviral activity, we employed enzymatic and chemical methods for selective glycan removal (horizontal cleavage), and a pool of proteases to cleave the peptide backbone (vertical cleavage), generating mucin fragments of varying size (**Figure 1b**). We then investigate the inhibition properties and binding to the whole virus of the horizontally and vertically cleaved mucin derivatives on IAV. We anticipate that, due to their higher abundance on mucins, sialylated *O*-glycans, are key factors for antiviral activity and we highlight a mucin size-dependent effect in viral inhibition.

## 2. Results and discussion

### 2.1. Purification and characterization of Bovine Submaxillary Mucin (BSM)

Bovine submaxillary mucin (BSM) is a commercially available mucin frequently used in biomaterial and biomedical applications (6, 13, 27). Due to the presence of sialylated glycans, it represents a suitable oral mucin model for interaction studies with pathogens, such as IAV. However, one of the major drawbacks of commercial mucins is their lack of purity. Due to their harsh processing, commercial mucins often contain numerous contaminants, such as endogenous proteases, small molecules, and processing additives, that can compromise their efficacy in various biological applications (28, 29). Aiming to minimize the effect of these contaminants, we purified BSM by size exclusion chromatography (**Figure S1a**) by modifying a protocol previously used for the purification of porcine gastric mucins (30).

A qualitative analysis of the purity of BSM after purification was checked by gel electrophoresis. To better appreciate variations in the high molecular weight region, we first conducted agarose gel electrophoresis. We observed that after purification, the band at an apparent molecular weight of ∼500 kDa, corresponding to either mucin fragments or glycosylated contaminants, was strongly reduced (**Figure S1b**). Differences in the smaller molecular weight region were visualized by SDS-PAGE. Here, multiple distinct bands below 100 kDa were visible in the unpurified product using both glycan and protein staining; on the contrary, after size exclusion chromatography purification, the same bands were completely or partially removed (**Figure 2a**). By that way, mucin content was increased, which became obvious from a lower protein and a higher carbohydrate content (**Figure S1c, d**). The higher carbohydrate content measured as PAS signal in samples at the same concentration of dry weight per volume points to an enrichment of the mucin fraction after purification. Distinct changes in composition between the crude and the purified BSM were revealed also by proteomic analysis (**Figure 2b**). Initially, 697 proteins were identified in the crude BSM; this number was reduced to around 660 proteins after purification (for complete list see Supplementary Data). Many of the remaining proteins (241) were significantly, and up to 16-times, reduced in their abundance after purification. The only type of mucin detected was the gel-forming MUC19 (**Figure 2c**), which shares similar domain organization and structural features with other secreted mucins such as MUC2, MUC5AC, MUC5b, and MUC6 (31, 32). Sequence alignment of bovine MUC19 (Uniprot P98091) with human MUC19 (Uniprot Q7Z5P9) reveals a 54% identity, highlighting substantial conservation between the two species. Notably, the proportion of MUC19 in the purified BSM product increased approximately threefold, from around 3.5% to about 11%. It is worth mentioning that for the quantification of MUC19 content we used the iBAQ algorithm to quantify absolute intensities. Even though the relative quantification is very robust with a large number of identified peptides, iBAQ values may underestimate absolute mucin levels as not all theoretical tryptic peptides can be measured. The highly glycosylated regions of mucins are highly resistant to proteolytic cleavage (vide infra), making detection of peptides in these regions challenging and thereby reducing the reported intensities.

**Figure 2.**
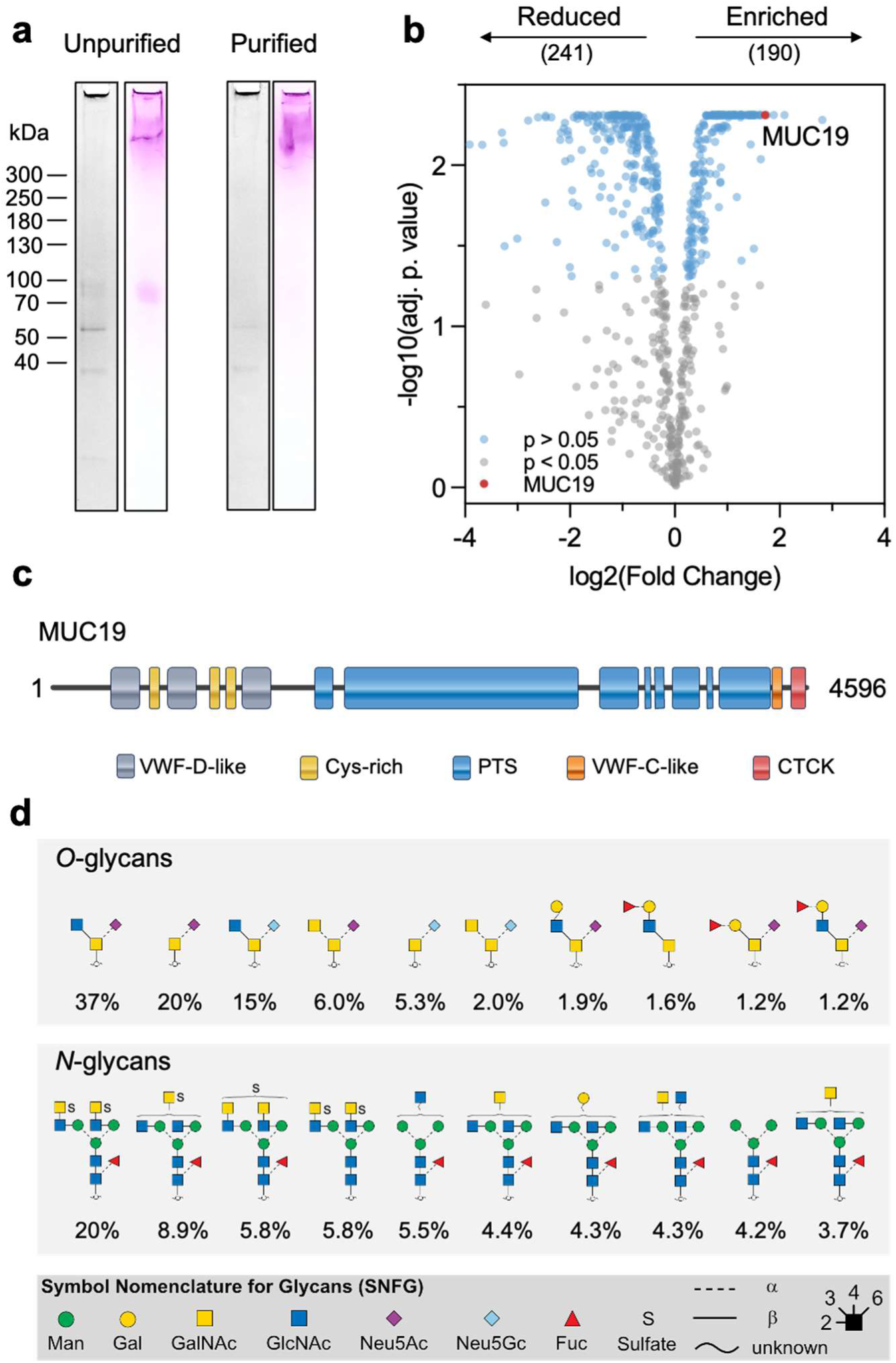
Purification of bovine submaxillary mucin (BSM) by size exclusion chromatography (SEC) enriches the mucin content. (**a**) SDS-PAGE stained with Coomassie blue (black and white) and periodic acid-Schiff (colored) of BSM before and after SEC purification. (**b**) Volcano plot of proteins with significantly different intensities (adjusted p-value <0.05) highlighted in blue. Mucin 19 (MUC19) is highlighted in red. On top of the plot are reported the number of proteins which abundance was reduced (241) and enriched (190) after purification. (**c**) Scheme of the structure of the gel forming bovine MUC19 based on the Uniprot entry P98091. (**d**) Putative structures and relative abundance of the top-10 of *O*- and *N*-glycans identified in the purified BSM.

*N*- and *O*-glycosylation of the purified BSM was analyzed by PGC-LC-MS/MS. A total of 40 *O*- and 24 *N*-glycans were identified (for complete assignment see Supplementary Data) and the top-10 most abundant structures are reported in **Figure 2d**. The top-10 most abundant *O*-glycan structures make up more than 90% of the BSM *O*-glycosylation. The identified *O*-glycans consisted of di- to heptasaccharides, however, more than 85% (expressed as relative abundance) of the *O*-glycans consist of small structures not larger than trisaccharides. As reported in previous studies, sialic acid units were found on 22 (93%) *O*-glycans, including 12 structures (69%) with Neu5Ac and 10 structures (24%) with Neu5Gc (25). The majority of the sialic acid units are α2-6 linked to the core GalNAc. Additionally, 20 structures (9%) carry one or more fucose units mostly at the terminal ends of the *O*-glycans. The majority of the *O*-glycan structures originate from core 3 (GlcNAcβ1-3GalNAc) with 57% and the Tn antigen (GalNAc) with 25%. The remaining *O*-glycans are based on core 1 (Galβ1-3GalNAc), 2 (Galβ1-3(GlcNAcβ1-6)GalNAc), 4 (GlcNAcβ1-3(GlcNAcβ1-6)GalNAc) and core 5 (GalNAcα1-3GalNAc). Among the identified *N*-glycans, 89% are complex-type, biantennary *N*-glycans. Similar to previous studies, we found that the *N*-glycans of BSM are low in sialic acids but high in sulfates and core fucoses (33, 34). Specifically, 11 structures (57%) are singly or doubly sulfated and 18 (83%) are fucosylated. Based on the glycan analysis, we conclude that both *N*- and *O*-glycosylation introduce a high density of negative charges to BSM. Besides their water-binding and gel-forming properties, sialylation, sulfation and fucosylation of the glycans may further provide multiple potential interaction sites for pathogens.

### 2.2. Glycan contribution to mucin’s antiviral activity against influenza

To investigate the structural determinants for influenza virus binding to mucins, mucin glycans from BSM were subjected to enzymatic or chemical removal of monosaccharides or specific glycans (**Table 1**, **Figure 3a**). Here, sialic acid was cleaved from purified BSM using a commercial mixture of sialidases (SialEXO^®^) with broad activity for α2-6, α2-3, and α2-8-linked sialic acids on both *O*- and *N*-glycans. Removal of *N*-glycans was achieved with peptide *N*-glycosidase F (PNGase-F), which is an amidase cleaving between the innermost GlcNAc and asparagine residues of almost all *N*-linked oligosaccharides. Since no universal enzyme for complete *O*-glycan removal from the heavily *O*-glycosylated regions of mucins has been identified so far, we chemically removed unsubstituted C3 GalNAc residues unspecifically by oxidative *β*-elimination following a protocol with minimal or no peptide core cleavage (**Figure S2**) (35).

**Figure 3.**
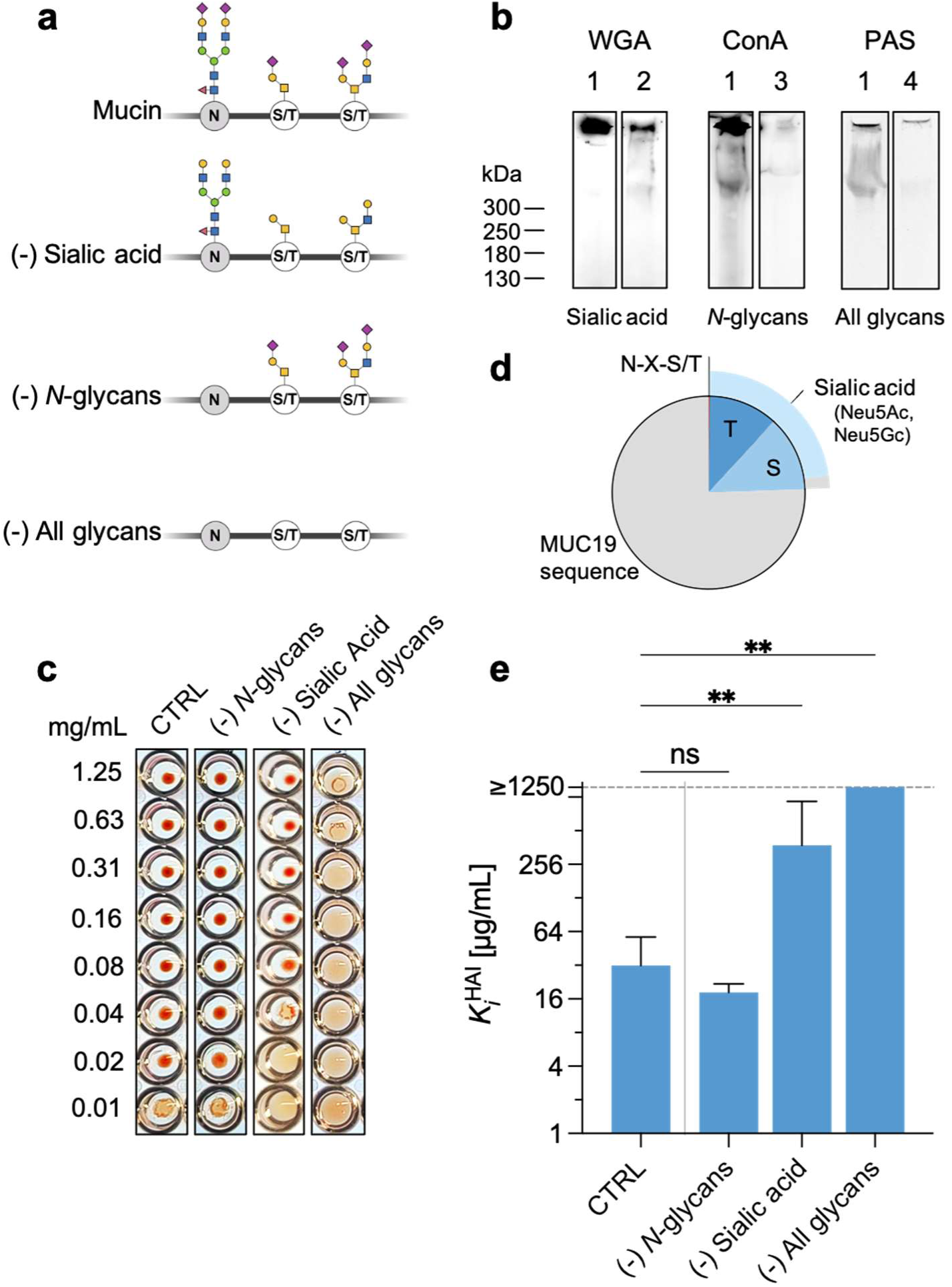
Investigation of the antiviral activity of mucin after selective removal of glycans. (**a**) Cartoon depicting the strategy pursued for selective removal of glycans from mucin. (**b**) SDS-PAGE / PAS and lectin blots for selective glycans staining (1 = untreated BSM, 2 = BSM after sialic acid removal, 3 = BSM after *N*-linked glycans removal, 4 = BSM after oxidation/β-elimination). (**c**) Hemagglutination inhibition assay using A/Panama/2007/1999(H3N2) virus (N≥3). (**d**) Pie chart with the abundance of predicted glycosylation on asparagine (N), threonine (T), and serine (S) in the bovine MUC19 (Uniprot entry P98091). The slice of the external pie chart reports the percentage of sialylated *O*-glycans according to the glycomic analysis of BSM. (**e**) Bar plot of the lowest log2 inhibitor concentration necessary to achieve complete inhibition of hemagglutination caused by the virus. No antiviral effect is observed for the sample without *O*- and *N*-glycans, therefore the K ^HAI^ was set over the maximum concentration of sample used in the experiment (*i.e.*, 1250 µg/mL). Intact purified BSM was used as control (CTRL). The significance for each group was calculated using the Kruskal-Wallis test. Results are displayed as the average (± SD) of N≥3 measurements. Results were compared by Dunn’s multiple comparison test. *p* < 0.05 (*), *p* < 0.01 (**), *p* < 0.001 (***), *p* < 0.0001 (****)

**Table 1:** Methods employed to selectively remove sialic acid or *O*- and *N*-linked glycans and their detection methods.

| Glycan type | Removal method | Detection method |
| --- | --- | --- |
| Sialic acid | Enzymatic – Sialidases derived from *Akkermansia muciniphila* | Lectin – Wheat germ agglutinin (WGA) |
| *N*-linked glycans | Enzymatic – PNGase-F | Lectin – Concanavalin A (ConA) |
| Unspecific (*O*- and *N*-linked glycans) | Chemical – Oxidation and β-elimination | Periodic acid – Schiff stain |

Product formation after glycan removal was analyzed using specific lectins for defined carbohydrates (**Figure 3b**). For sialic acid, we used fluorescently labelled wheat germ agglutinin (WGA) to follow the efficiency of carbohydrate removal on BSM (36, 37), while to visualize *N*-linked glycans, we used fluorescently labelled high-mannose-binding concanavalin A (Con-A), as Con-A specifically binds to *N*-glycans due to their high mannose content (38, 39). Periodic acid-Schiff stain (PAS) was used as a universal method to visualize the glycans following the unspecific release of mucin glycans through *β*-elimination treatment (40). The SDS-PAGE showed that PNGase-F treatment effectively removed *N*-glycans, while *β*-elimination led to the removal of nearly all glycans. In contrast, WGA staining of sialic acid revealed a reduced but still present sialylated sugars even after treatment with sialidases. Further, quantitative sialic acid measurements (NANA assay) showed that the sialidases cleaved approximately 67% (**Figure S3**) of the total sialic acid on BSM, in agreement with previous findings, suggesting the presence of sialic acid residues that may be inaccessible or resistant to cleavage (41).

Next, the potential of BSM or deglycosylated derivatives of BSM to prevent binding of seasonal influenza virus (A/Panama/2007/99 (H3N2)) to human red blood cells (hRBC) was investigated by the well-established hemagglutination inhibition (HAI) assay (**Figure 3c**). The inhibition effectiveness was expressed as K*_i_*^HAI^ representing the lowest concentration of inhibitor that successfully inhibits hemagglutination at four hemagglutination units (4 HAU). BSM was used as a control, showing the highest inhibition constant (K*_i_*^HAI^) of about 30 µg/mL (21.3 nM) (**Figure 3e**). Interestingly, even though the purification of BSM increased its overall quality by enriching the mucin content and lowering the contaminants, this was not translated also in a variation of its antiviral activity as measured by the HAI assay. This observation suggests that protein contaminants which are present in BSM do not, or minimally interfere with the binding of mucins to the virus.

We found that partially removing sialic acid from BSM significantly decreases by over 10-fold but does not eliminate antiviral inhibitory activity (K*_i_*^HAI^ = 380 µg/mL). Since sialic acid was not completely cleaved by sialidase treatment, this might explain the observed remnant activity. Removal of *N*-glycans does not significantly affect the antiviral activity of BSM. This observation could be related to different factors; firstly, *N*-linked glycans are comparatively less abundant than *O*-linked glycans. Glycosylation predictions using NetNGlyc (42) indicate that less than 1% of the asparagine residues as part of the N-X-S/T sequon in the MUC19 sequence are glycosylated; in contrast, predictions from NetOGlyc (43) reveals that approximately 24% of serine and threonine residues are *O*-glycosylated (**Figure 3d)**. Secondly, the glycomic analysis highlighted that the analyzed *N*-glycans from BSM are low in sialic acids, which we confirmed to be pivotal for influenza inhibition. The low abundancy and scarce sialylation might explain the negligible participation of *N*-glycans in virus interaction. Previous studies proposed *N*-glycosylation as key feature in mucin synthesis, stability, and folding rather than direct interaction with viruses (44, 45). Although the *N*-glycans of mucin might have a negligible effect on the antiviral activity of isolated mucin, it is important to note that mucus consists also of other non-mucin proteins (*e.g.*, lysozyme, lactoferrin, surfactant proteins). Other *N*-glycosylated non-mucin proteins may interact with the virus in ways that are underestimated in isolated BSM.

On the other hand, removing all glycans unspecifically, resulted in abolished antiviral inhibitory activity, with no inhibition observed even at the highest concentration tested (K*_i_*^HAI^ > 1250 µg/mL), suggesting a predominant role of mucin *O*-glycans in influenza virus engagement. It is worth mentioning that because of the unspecificity of the *β*-elimination reaction, removal of *O*-glycans implies the simultaneous removal of sialic acid, accounting then for the combined effect of *O*-glycans and sialic acid. Nevertheless, these findings point at sialylated *O*-glycans as the primary interaction sites for influenza virus, while exclude *N*-glycans from playing a significant role in viral binding.

### 2.3. Mucin size correlates with antiviral effect

The size of polymeric antiviral materials can impact its efficacy in binding to the virus (22, 46), therefore, we investigated how the size of the mucin polymers influences its antiviral effect. Because of their polymeric structure and high glycosylation, mucins exhibit intrinsic resistance to proteolytic degradation. Complete degradation requires an arsenal of carbohydrate-active enzymes and proteases able to degrade the glycans and the peptide core (47, 48). To obtain mucin glycosylated fragments of different sizes, we digested BSM using proteases with different cleavage specificities (**Table 2**). We selected a diverse set of proteolytic enzymes, including animal and plant proteases as well as mucinases, and conducted the reactions under optimal conditions specific to each enzyme’s activity. After overnight digestion, cleavage efficiency was assessed by SDS-PAGE and PAS staining, while the antiviral activity of the fragmented BSM was measured by hemagglutination inhibition.

**Table 2:** Specificity and origin of the proteases used to cleave BSM.

| Protease | Family | Specificity | Reference |
| --- | --- | --- | --- |
| SmE | Metalloprotease | N-terminally to a glycosylated Ser or Thr residue.<br>(Different O-glycans at the P1' position are accepted). | (50) |
| StcE | Metalloprotease | N-terminally to a Ser/Thr*-X-Ser/Thr, where the cleavage happens on the amino side of the 2<sup>nd</sup> Ser/Thr, and X can be any amino acid, or none. | (51) |
| Pepsin | Aspartic protease | Broad specificity: preference for cleavage C-terminal to Phe, Leu, and Glu. | (52) |
| Chymotrypsin | Serine protease | C-terminal of hydrophobic residues (*e.g.*, Phe, Tyr, Trp, Leu). | (52) |
| Trypsin | Serine protease | C-terminal of Lys and Arg residues, except for -Arg-Pro- and -Lys-Pro- bonds which are normally resistant to proteolysis. | (52)·(53) |
| Elastase | Serine protease | C-terminal of amino acids with small hydrophobic side chains (e.g., Ala, Val, Ser, Gly, Leu, Ile) | (52) |
| Proteinase K | Serine protease | Broad specificity: C-terminal of aliphatic and aromatic amino acids. | (52, 54) |
| Bromelain | Cysteine endopeptidase | Broad specificity: cleaves at the C-terminal, high efficiency on Arg-Arg-containing substrates. | (55) |
| Papain | Cysteine endopeptidase | Broad specificity: cleaves at the C-terminal, preference for amino acids bearing a large hydrophobic side chain at the P2 position. Does not accept Val in P1'. | (56) |
| Ficin (ficain) | Cysteine endopeptidase | Broad specificity: cleaves at the C-terminal, accepts Gly, Ser, Thr, Met, Lys, Arg, Tyr, Ala, Asn, and Val, with preferences for hydrophobic side chains. | (57) |

Intact purified BSM becomes visible by PAS after agarose gel electrophoresis as a broad band in the MDa range, with an estimated apparent molecular weight of about 1.5 MDa (**Figure S1b**) (49). Except for *Serratia marcescens* Enhancin (SmE), the digestion of BSM with all the proteases displayed a mass distribution of the fragments at lower molecular mass range compared to the intact BSM (**Figure 4a**). Among the tested proteases, proteinase K demonstrated the highest efficiency in degrading mucin, producing glycosylated fragments ranging from approximately 40 to 70 kDa. Fragments in such a size range could be indicative of the smallest structural tandem repeat units that retain extensive glycosylation and are resistant to proteolytic cleavage. Antiviral activity assessed by hemagglutination inhibition was observed only for the BSM digests obtained with SmE, StcE, and ficin at a concentration up to 1250 µg/mL (**Figure 4b, c**). In contrast, digests from all the other enzymes, including pepsin, chymotrypsin, trypsin, elastase, bromelain, papain, and proteinase K, exhibited no visible antiviral activity (*i.e.*, K*_i_*^HAI^ values exceeding 1250 µg/mL) (**Figure 4c**). Based on the mass distribution of the BSM fragments observed on the SDS-PAGE gel, a discernible size-activity relationship can be inferred. Specifically, a trend emerges indicating that the antiviral activity of the glycosylated fragments is dependent on their size (**Figure 4d**), specifically as the molecular weight increases, the K*_i_*^HAI^ decreases in a non-linear manner, suggesting that larger structures exhibit higher antiviral activity. Notably, rough thresholds can be traced around 850 and 200 kDa defining regions where the size-dependent activity drops by a factor of 10x and 100x.

**Figure 4.**
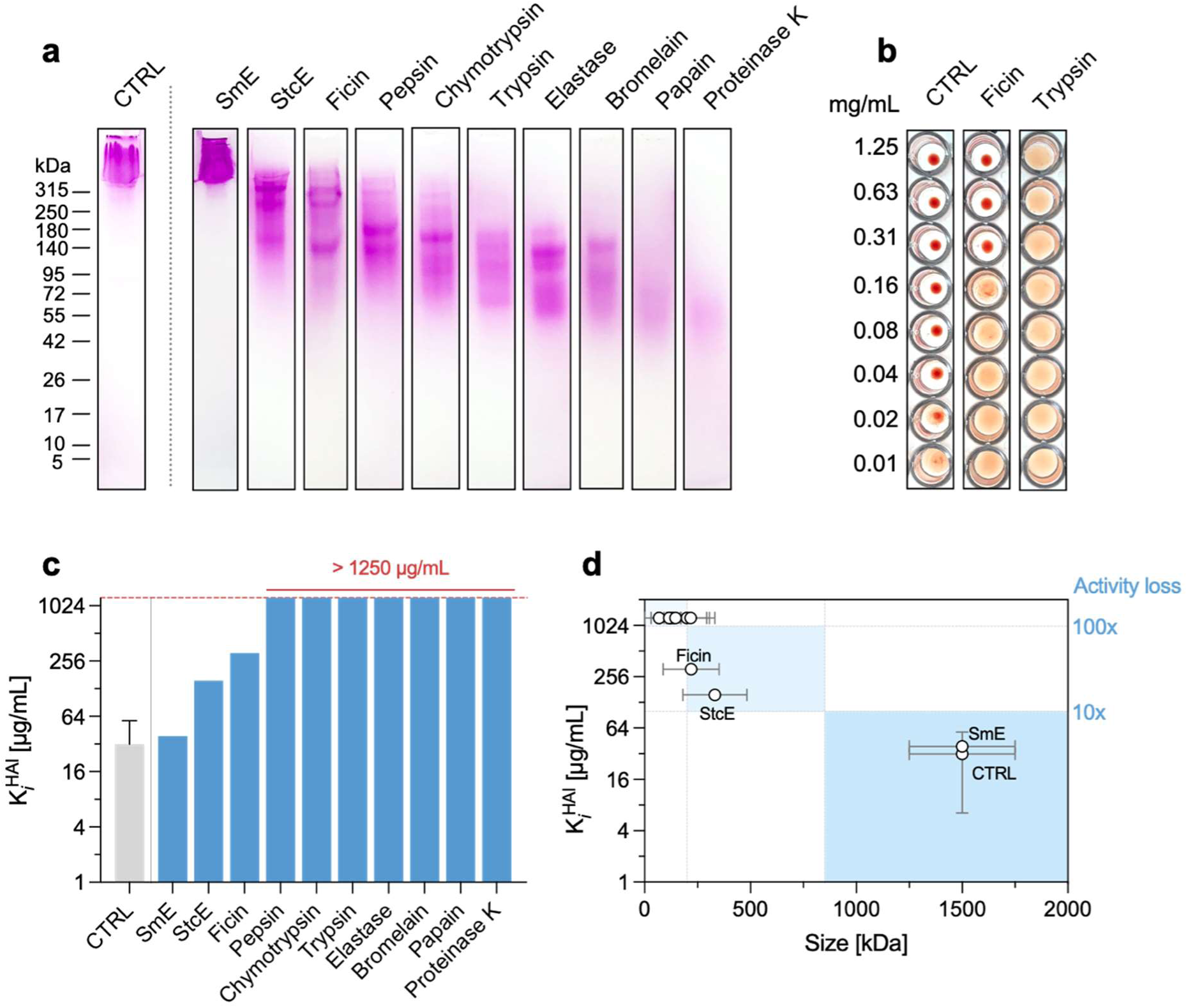
Quantifying the effect of the size of mucins on the inhibition activity on virus induced hemagglutination. Bovine submaxillary mucin (BSM) was digested with specific and unspecific proteases to obtain mucin fragments of different sizes. (**a**) Cleavage effectiveness was assessed by SDS-PAGE and PAS staining. (**b**) The antiviral activity of the fragmented BSM was assessed by hemagglutination inhibition assay (reported only the control sample (PBS), ficin, and trypsin as representative of active, partially, and inactive samples, respectively). (**c**) Inhibition constant (K*_i_*^HAI^) defined as the lowest concentration of BSM or BSM fragments that is able to inhibit agglutination. Results are displayed as the average (± SD, for some samples not visible as being too small) of N≥3 measurements. (**d**) Relationship between K_i_ ^HAI^ and the size of mucin or mucin fragments with activity regions highlighted in blue. Arbitrary thresholds at about 850 and 200 kDa have been set to delineate regions where the activity undergoes reduction by one and two orders of magnitude.

### 2.4. Glycosylated mucin fragments retain the virus-binding capacity and antiviral properties in a size-dependent manner

The hemagglutination inhibition assay of BSM digests indicates that the size of the mucin fragments (MFs) influences their antiviral efficacy. To explore this relationship further, we investigated how the molecular size of these fragments impacts their binding affinity to the influenza virus. We enriched and purified glycosylated MFs with varying molecular weights and different antiviral activities. Specifically, we selected glycosylated BSM fragments generated by StcE (MF1) and proteinase K (MF2) digestions, representing the largest and smallest fragments detectable by SDS-PAGE in our experimental conditions (**Figure 5a**). Compared to native BSM, which exhibits an apparent molecular weight in the MDa range (**Figure S1b**), MF1 and MF2 are revealed by PAS at ∼330 kDa and ∼50 kDa, respectively. Additionally, the hydrodynamic radii of these fragments differ by several order of magnitudes, with intact BSM measuring over 2000 nm, while for MF1 and MF2 measuring 60 nm and 2.5 nm, respectively (**Figure 5b**). In terms of sialic acid composition, both fragments have less abundancy compared to BSM. Specifically, BSM contains over 550 mol of SA per mol of protein, while MF1 and MF2 contain approximately 390 and 26 mol of SA per mol of protein, respectively. When normalized by the apparent molecular weight to express the density of SA, MF1 shows the highest density at 1.2 mol/kDa, surpassing both BSM (0.4 mol/kDa) and MF2 (0.5 mol/kDa) (**Figure S4**). This suggests that both MFs retain a high sialic acid density. Structurally these fragments could mainly, if not only, be constituted of the highly glycosylated regions (*i.e.*, PTS domains) of mucin, with minimum or any non-glycosylated parts.

**Figure 5.**
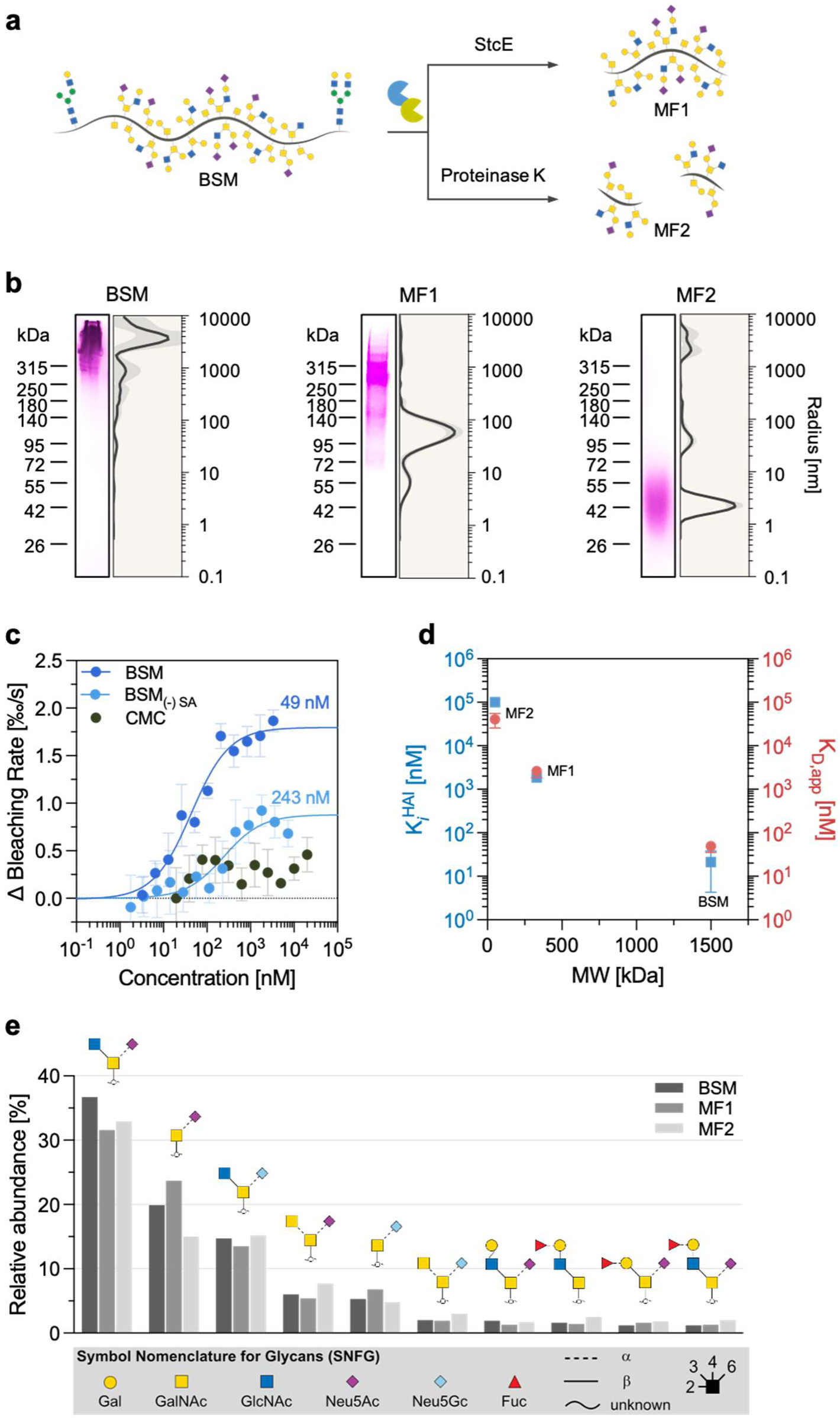
Mucin fragments have virus-binding and antiviral capacity in a size-dependent manner. (**a**) Illustration of the digestion of mucin from the bovine submaxillary gland (BSM) with StcE (MF1) or proteinase K (MF2). (**b**) SDS-PAGE/PAS staining and dynamic light scattering of the native BSM, and the purified mucin fragments. (**c**) Microscale thermophoresis (MST) change in the bleaching rate upon binding of BSM at different concentrations to fluorescently labelled R18 A/X31 virus at the steady state. Desialylated BSM (BSM_(-)SA_) and carboxymethyl cellulose (CMC) served as controls. Binding constants are described as apparent dissociation constants (K_D,app_). Each data point represents the average values of N=4 experiments and the error bars show the standard deviation. Data points were fitted according to the mass-action law function to calculate K_D,app_ values. (**d**) HAI assay against four hemagglutination units (HAU) A/X31 and the K_D,app_ measured by MST. (**e**) Relative abundance and putative structures of the top-10 *O*-glycans in BSM, MF1, and MF2.

The HAI assay revealed that MF1 and MF2 have HAI constants (K*_i_*^HAI^) of 1.88 µM (0.63 mg/mL) and 100 µM (5 mg/mL), respectively (**Figure S5 a, c**). When compared to BSM, these values reflect a reduction of antiviral activity by approximately two and four orders of magnitude for MF1 and MF2, respectively. Previous studies with sialylated bioinspired brush polymers (∼400 kDa) demonstrated inhibition of influenza virus (H1N1 subtype) with a K*_i_*^HAI^ of 0.24 µM, placing its activity in a comparable range to MF1 (58).

Next, we characterized the affinity at which the different BSM derivatives bind to influenza A/X31 virus (H3N2 subtype), which is a subtype antigenically similar to A/Panama/2007/99 but with differences in the glycosylation sites (59, 60). Microscale thermophoresis (MST) experiments showed that the intact BSM had an apparent dissociation constant (K_D,app_) in the nanomolar range (49 nM) (**Figure 5c**). Desialylated BSM (BSM_(-)SA_) was used as binding control (*i.e.*, expected lower binding than BSM), while carboxymethyl cellulose (CMC), a high molecular weight natural polymer similar to BSM in viscosity and charge, served as a negative control binder. Partial cleavage of sialic acid (∼67% efficacy, **Figure S3**) from BSM reduced its binding affinity for the virus by approximately 5-fold (243 nM). This suggests that while reduced sialylation in BSM still permits virus binding, it does so with reduced effectiveness, as indicated by the lower affinity and binding amplitude compared to intact BSM.

MST measurements with MF1 and MF2 (**Figure S5 b, d**) supported the HAI assay findings, showing very good agreement between K_D,app_ and K*_i_*^HAI^ values (**Figure S6**). The reduction in fragment size weakened both the binding avidity and the antiviral inhibitory capacity, evidenced by an exponential increase in both K_D,app_ and K*_i_*^HAI^ (**Figure 5d**). These observations are consistent with the notion that larger glycoprotein structures can present multiple binding sites, thereby enhancing overall viral binding and inhibitory activity (22, 58, 61, 62). The ability of the MFs to bind to the virus suggests that the glycan structures involved in virus attachment are still present in the small fragments. *O*-glycan analysis showed that the same glycans identified in intact BSM are still present, and with comparable relative abundance, on MF1 and MF2 (**Figure 5e**). Interestingly, assuming the MFs as individual (MF2) or serial repetition (MF1) of tandem repeats of the highly glycosylated regions (PTS) of MUC19, we can conclude that the glycosylation pattern is conserved and homogeneously distributed on the tandem repeats of mucin. Yet, the reduced size and reduced multiplicity of binding sites results in weaker and less effective inhibition. Previous studies have proposed that an ideal antiviral polymer should position sialic acid residues at intervals around the intra-hemagglutinin spacing (∼4.5 nm) while maintaining an overall polymer length larger than the inter-hemagglutinin trimer spacing (∼14 nm) (58, 63). This configuration would ensure a high local concentration of sialic acids and the ability to bridge multiple hemagglutinin trimers, thereby enhancing multivalent binding and viral inhibition. Based on these criteria, MF2 (∼2.5 nm radius) alone as an antiviral could roughly span the intra-hemagglutinin distance giving place to bivalent binding, but it fails to meet the necessary spatial arrangement for inter-hemagglutinin binding, limiting its capacity for effective viral engagement. Similarly, although MF1 could potentially meet both criteria due to its larger size (∼60 nm radius) and high SA abundance, its antiviral activity may still be insufficient on its own.

Rather than functioning independently, mucin fragments could serve as versatile building blocks for larger antiviral architectures (*e.g.*, linear polymers, polymeric networks), leveraging both their glycan diversity and steric effects. Importantly, these mucin-derived fragments, as natural glycopeptides, present minimal functional virus-binding units with naturally evolved glycan diversity and optimized spatial distribution. Unlike biomimetic polymers that require meticulous design to mimic these features, mucin-derived glycopeptides inherently combine structural simplicity with functional efficacy. This dual advantage positions them as promising candidates for the development of novel broad-spectrum antiviral biomaterials, where the combination of steric contribution and multiple weak binding interactions can significantly amplify inhibitory effects.

## 3. Conclusions

In summary, we explored how structural features of bovine submaxillary mucin influence its antiviral activity against influenza virus (H3N2 subtype). We highlighted the crucial role of sialylated *O*-glycans in defining the antiviral activity of BSM, with *N*-glycans playing a minimal role in virus interaction, possibly because of the low level of sialylation. While partial removal of sialic acid reduced but did not completely abolish antiviral activity, full glycan removal eliminated inhibition altogether, underscoring the importance of *O*-glycans. We showed that mucin fragments of different size displaying the same *O*-glycosylation pattern as full-length mucin, can be obtained by proteolytic digestion and we observed a strong size-dependent effect on antiviral efficacy. Intact BSM (MDa scale) exhibited high viral binding affinity in the nanomolar range and potent inhibition, while smaller mucin fragments (MF1, ∼330 kDa, and MF2, ∼50 kDa) showed significantly reduced antiviral activity with K*_i_*^HAI^ values of 0.63 mg/mL and 5 mg/mL, respectively. Binding affinity decreased proportionally, with K_D,app_ values increasing to 1.88 µM for MF1 and 100 µM for MF2.

These results emphasize that larger glycoprotein structures, with their higher glycan density and multiple binding sites, enhance both virus binding and inhibition. This size-dependent antiviral response highlights the importance of mucin polymer length in mediating effective viral neutralization. Moreover, the ability of small mucin fragments, such as MF2 to bind influenza virus, points to potential applications as viral decoys for biopolymers or nanoparticles for targeted drug delivery. Further engineering of hydrogels or nanoparticles harboring such building blocks can lead to novel antiviral materials with economic use of antiviral ligands arranged in nanoclusters.

## 4. Materials and Methods

Full details on materials and methods are described in SI Appendix. The methods, include mucin purification with proteomic and glycomic analysis; chemical and enzymatic methods used to remove glycans from purified BSM; enzymatic mucin fragmentation and purification of glycosylated mucin fragments; expression and purification of mucinases; gel electrophoresis and blotting; hemagglutination assay; and microscale thermophoresis.

## Supporting information

Supplementary_Data_Protein_List

Supplementary_Data_Glycan_List

## Author Contributions

Conception and design of the study: C.B., D.L. Acquisition, analysis, and interpretation of the data: C.B., J.T., T.L.P., R.D., K.F., M.S. Drafting the article or revising it critically for important intellectual content: C.B., J.T., T.L.P., R.D., K.F., M.S., P.M., K.P., D.L. All authors have given approval to the final version of the manuscript.

## Acknowledgements

We are grateful to Stacy Malaker for providing us with pet28a_SmE that we used to express SmE, Junqioa Jia in the Markus Wahl lab (Freie Universität Berlin) for support in expression and purification of StcE, and professor Dr. Joachim Heberle for giving us access to his Monolith instrument to perform MST studies. The project was realized with research infrastructure and support provided by the Research Building SupraFAB realized with funds from the Federal Government (BMBF) and the State of Berlin. DL is grateful for financial support from the federal ministry of education and research funded project MucPep (FKZ: 13XP511). The authors acknowledge the financial support of the Collaborative Research Center “Dynamic Hydrogels at Biological Interfaces” (CRC 1449) funded by the Deutsche Forschungsgemeinschaft (DFG, German Research Foundation) – Project ID 431232613– SFB 1449 (sub-projects C03, Z01 and the IRTG).

## References

1. L. Mackenzie, World Health Organization - The burden of Influenza. (2024). Available at: https://www.who.int/news-room/feature-stories/detail/the-burden-of-influenza#:~:text=Disease burden from seasonal influenza&text=Influenza%2C or the flu%2C is,viruses%2C after the common cold.

2. S. Malik, M. Asghar, Y. Waheed, Outlining recent updates on influenza therapeutics and vaccines: A comprehensive review. Vaccine X 17, 100452 (2024).

3. L. Jiang, H. Chen, C. Li, Advances in deciphering the interactions between viral proteins of influenza A virus and host cellular proteins. Cell Insight 2, 100079 (2023).

4. C. L. Wardzala, A. M. Wood, D. M. Belnap, J. R. Kramer, Mucins Inhibit Coronavirus Infection in a Glycan-Dependent Manner. ACS Cent. Sci. 8, 351–360 (2022).

5. I. Ethan, et al., Membrane-Tethered Mucin 1 Is Stimulated by Interferon and Virus Infection in Multiple Cell Types and Inhibits Influenza A Virus Infection in Human Airway Epithelium. MBio 13, e01055–22 (2022).

6. M. Kretschmer, et al., Synthetic Mucin Gels with Self-Healing Properties Augment Lubricity and Inhibit HIV-1 and HSV-2 Transmission. Adv. Sci. 9, 2203898 (2022).

7. R. Bansil, B. S. Turner, Mucin structure, aggregation, physiological functions and biomedical applications. Curr. Opin. Colloid Interface Sci. 11, 164–170 (2006).

8. M. Cohen, et al., Influenza A penetrates host mucus by cleaving sialic acids with neuraminidase. Virol. J. 10, 321 (2013).

9. J. N. S. S. Couceiro, J. C. Paulson, L. G. Baum, Influenza virus strains selectively recognize sialyloligosaccharides on human respiratory epithelium; the role of the host cell in selection of hemagglutinin receptor specificity. Virus Res. 29, 155–165 (1993).

10. J. Mayr, et al., Unravelling the Role of O-glycans in Influenza A Virus Infection. Sci. Rep. 8, 16382 (2018).

11. C. M. Spruit, et al., N-Glycolylneuraminic Acid in Animal Models for Human Influenza A Virus. Viruses 13 (2021).

12. T. Takahashi, et al., *N*-Glycolylneuraminic Acid on Human Epithelial Cells Prevents Entry of Influenza A Viruses That Possess *N*-Glycolylneuraminic Acid Binding Ability. J. Virol. 88, 8445–8456 (2014).

13. L. Kaler, et al., Influenza A virus diffusion through mucus gel networks. Commun. Biol. 5, 249 (2022).

14. K. Barnard, et al., Modified Sialic Acids on Mucus and Erythrocytes Inhibit Influenza A Virus Hemagglutinin and Neuraminidase Functions. J. Virol. 94, 10.1128/jvi.01567-19 (2020).

15. G. Petrou, T. Crouzier, Mucins as multifunctional building blocks of biomaterials. Biomater. Sci. 6, 2282–2297 (2018).

16. G. W. Hughes, et al., The MUC5B mucin polymer is dominated by repeating structural motifs and its topology is regulated by calcium and pH. Sci. Rep. 9, 17350 (2019).

17. R. W. H. Ruigrok, L. J. Calder, S. A. Wharton, Electron microscopy of the influenza virus submembranal structure. Virology 173, 311–316 (1989).

18. S. Bhatia, et al., Linear polysialoside outperforms dendritic analogs for inhibition of influenza virus infection in vitro and in vivo. Biomaterials 138, 22–34 (2017).

19. C. S. Delaveris, E. R. Webster, S. M. Banik, S. G. Boxer, C. R. Bertozzi, Membrane-tethered mucin-like polypeptides sterically inhibit binding and slow fusion kinetics of influenza A virus. Proc. Natl. Acad. Sci. 117, 12643–12650 (2020).

20. W. J. Lees, A. Spaltenstein, J. E. Kingery-Wood, G. M. Whitesides, Polyacrylamides Bearing Pendant .alpha.-Sialoside Groups Strongly Inhibit Agglutination of Erythrocytes by Influenza A Virus: Multivalency and Steric Stabilization of Particulate Biological Systems. J. Med. Chem. 37, 3419–3433 (1994).

21. S.-K. Choi, M. Mammen, G. M. Whitesides, Generation and in Situ Evaluation of Libraries of Poly(acrylic acid) Presenting Sialosides as Side Chains as Polyvalent Inhibitors of Influenza-Mediated Hemagglutination. J. Am. Chem. Soc. 119, 4103–4111 (1997).

22. M. Wallert, et al., Mucin-Inspired, High Molecular Weight Virus Binding Inhibitors Show Biphasic Binding Behavior to Influenza A Viruses. Small 16, 2004635 (2020).

23. V. R. Kohout, C. L. Wardzala, J. R. Kramer, Synthesis and biomedical applications of mucin mimic materials. Adv. Drug Deliv. Rev. 191, 114540 (2022).

24. T. Jaroentomeechai, et al., Mammalian cell-based production of glycans, glycopeptides and glycomodules. Nat. Commun. 15, 9668 (2024).

25. M. Safferthal, L. Bechtella, A. Zappe, G. M. Vos, K. Pagel, Labeling of Mucin-Type O-Glycans for Quantification Using Liquid Chromatography and Fluorescence Detection. ACS Meas. Sci. Au 4, 223–230 (2024).

26. J. Kim, et al., Structural and Quantitative Characterization of Mucin-Type O-Glycans and the Identification of O-Glycosylation Sites in Bovine Submaxillary Mucin. Biomolecules (2020).

27. H. Rulff, et al., Comprehensive Characterization of the Viscoelastic Properties of Bovine Submaxillary Mucin (BSM) Hydrogels and the Effect of Additives. Biomacromolecules (2024). 10.1021/acs.biomac.4c00153.

28. G. Tettamanti, W. Pigman, Purification and characterization of bovine and ovine submaxillary mucins. Arch. Biochem. Biophys. 124, 41–50 (1968).

29. M. Marczynski, C. Kimna, O. Lieleg, Purified mucins in drug delivery research. Adv. Drug Deliv. Rev. 178, 113845 (2021).

30. V. J. Schömig, et al., An optimized purification process for porcine gastric mucin with preservation of its native functional properties. RSC Adv. 6, 44932–44943 (2016).

31. Y. Chen, et al., Genome-Wide Search and Identification of a Novel Gel-Forming Mucin MUC19/Muc19 in Glandular Tissues. Am. J. Respir. Cell Mol. Biol. 30, 155–165 (2004).

32. D. Fass, D. J. Thornton, Mucin networks: Dynamic structural assemblies controlling mucus function. Curr. Opin. Struct. Biol. 79, 102524 (2023).

33. J. Kim, et al., N-Glycan Modifications with Negative Charge in a Natural Polymer Mucin from Bovine Submaxillary Glands, and Their Structural Role. Polym. (2021).

34. J. Kim, et al., N-glycans of bovine submaxillary mucin contain core-fucosylated and sulfated glycans but not sialylated glycans. Int. J. Biol. Macromol. 138, 1072–1078 (2019).

35. T. A. Gerken, R. Gupta, N. Jentoft, A novel approach for chemically deglycosylating O-linked glycoproteins. The deglycosylation of submaxillary and respiratory mucins. Biochemistry 31, 639–648 (1992).

36. V. P. Bhavanandan, A. W. Katlic, The interaction of wheat germ agglutinin with sialoglycoproteins. The role of sialic acid. J. Biol. Chem. 254, 4000–4008 (1979).

37. B. P. Peters, S. Ebisu, I. J. Goldstein, M. Flashner, Interaction of wheat germ agglutinin with sialic acid. Biochemistry 18, 5505–5511 (1979).

38. C. F. Brewer, Interactions of an ovalbumin glycopeptide with concanavalin A. Biochem. Biophys. Res. Commun. 90, 117–122 (1979).

39. D. Bojar, et al., A Useful Guide to Lectin Binding: Machine-Learning Directed Annotation of 57 Unique Lectin Specificities. ACS Chem. Biol. 17, 2993–3012 (2022).

40. M. Mantle, A. Allen, A colorimetric assay for glycoproteins based on the periodic acid/Schiff stain [proceedings]. Biochem. Soc. Trans. 6, 607–609 (1978).

41. H. Yan, et al., Glyco-Modification of Mucin Hydrogels to Investigate Their Immune Activity. ACS Appl. Mater. Interfaces 12, 19324–19336 (2020).

42. R. Gupta, S. Brunak, Prediction of glycosylation across the human proteome and the correlation to protein function. Pac. Symp. Biocomput. 310–322 (2002).

43. C. Steentoft, et al., Precision mapping of the human O-GalNAc glycoproteome through SimpleCell technology. EMBO J. 32, 1478–1488 (2013).

44. T. Taniguchi, et al., N-Glycosylation affects the stability and barrier function of the MUC16 mucin. J. Biol. Chem. 292, 11079–11090 (2017).

45. S. Parry, et al., N-Glycosylation of the MUC1 mucin in epithelial cells and secretions. Glycobiology 16, 623–634 (2006).

46. J. Vonnemann, et al., Size Dependence of Steric Shielding and Multivalency Effects for Globular Binding Inhibitors. J. Am. Chem. Soc. 137, 2572–2579 (2015).

47. J. F. Wardman, R. K. Bains, P. Rahfeld, S. G. Withers, Carbohydrate-active enzymes (CAZymes) in the gut microbiome. Nat. Rev. Microbiol. 20, 542–556 (2022).

48. C. Wickström, M. C. Herzberg, D. Beighton, G. Svensäter, Proteolytic degradation of human salivary MUC5B by dental biofilms. Microbiology 155, 2866–2872 (2009).

49. L. Shi, K. D. Caldwell, Mucin Adsorption to Hydrophobic Surfaces. J. Colloid Interface Sci. 224, 372–381 (2000).

50. J. Chongsaritsinsuk, et al., Glycoproteomic landscape and structural dynamics of TIM family immune checkpoints enabled by mucinase SmE. Nat. Commun. 14, 6169 (2023).

51. S. A. Malaker, et al., The mucin-selective protease StcE enables molecular and functional analysis of human cancer-associated mucins. Proc. Natl. Acad. Sci. 116, 7278–7287 (2019).

52. P. J. Sweeney, J. M. Walker, “Proteinase K (EC 3.4.21.14) BT - Enzymes of Molecular Biology” in M. M. Burrell, Ed. (Humana Press, 1993), pp. 305–311.

53. M. Manea, G. Mező, F. Hudecz, M. Przybylski, Mass spectrometric identification of the trypsin cleavage pathway in lysyl-proline containing oligotuftsin peptides. J. Pept. Sci. 13, 227–236 (2007).

54. W. Ebeling, et al., Proteinase K from Tritirachium album Limber. Eur. J. Biochem. 47, 91–97 (1974).

55. D. N. Gosalia, C. M. Salisbury, J. A. Ellman, S. L. Diamond, High Throughput Substrate Specificity Profiling of Serine and Cysteine Proteases Using Solution-phase Fluorogenic Peptide Microarrays*. Mol. Cell. Proteomics 4, 626–636 (2005).

56. A. C. Storer, R. Ménard, “Chapter 419 - Papain” in Handbook of Proteolytic Enzymes, N. D. Rawlings, G. B. T.-H. of P. E. (Third E. Salvesen, Eds. (Academic Press, 2013), pp. 1858–1861.

57. A. Singleton, D. J. Buttle, “Chapter 428 - Ficain” in Handbook of Proteolytic Enzymes, N. D. Rawlings, G. B. T.-H. of P. E. (Third E. Salvesen, Eds. (Academic Press, 2013), pp. 1877–1879.

58. S. Tang, et al., Antiviral Agents from Multivalent Presentation of Sialyl Oligosaccharides on Brush Polymers. ACS Macro Lett. 5, 413–418 (2016).

59. S. C. B. Gopinath, et al., An RNA aptamer that distinguishes between closely related human influenza viruses and inhibits haemagglutinin-mediated membrane fusion. J. Gen. Virol. 87, 479–487 (2006).

60. M. D. Tate, et al., Playing Hide and Seek: How Glycosylation of the Influenza Virus Hemagglutinin Can Modulate the Immune Response to Infection. Viruses 6, 1294–1316 (2014).

61. G. B. Sigal, M. Mammen, G. Dahmann, G. M. Whitesides, Polyacrylamides Bearing Pendant α-Sialoside Groups Strongly Inhibit Agglutination of Erythrocytes by Influenza Virus: The Strong Inhibition Reflects Enhanced Binding through Cooperative Polyvalent Interactions. J. Am. Chem. Soc. 118, 3789–3800 (1996).

62. M. Nagao, et al., Design of Glycopolymers Carrying Sialyl Oligosaccharides for Controlling the Interaction with the Influenza Virus. Biomacromolecules 18, 4385–4392 (2017).

63. V. Bandlow, et al., Sialyl-LacNAc-PNA⋅DNA Concatamers by Rolling-Circle Amplification as Multivalent Inhibitors of Influenza A Virus Particles. ChemBioChem 20, 159–165 (2019).

