## Supplementary Information for "Structural Determinants of Mucins in Influenza Virus Inhibition: The Synergistic Role of Sialylated Glycans and Molecular Size"

**Figures**

- **Figure S1**: Purification of mucins from bovine submaxillary gland (BSM) by size exclusion chromatography and comparison with crude BSM.
- **Figure S2**: SDS-PAGE of BSM before and after oxidative $\beta$-elimination.
- **Figure S3**: Sialic acid quantification after sialidase treatment of BSM.
- **Figure S4**: Sialic acid abundance and density in mucin from bovine submaxillary mucin (BSM) and in mucin fragments.
- **Figure S5**: Antiviral activity and binding to influenza virus of mucin fragments.
- **Figure S6**: Correlation plot between virus inhibition and virus binding.

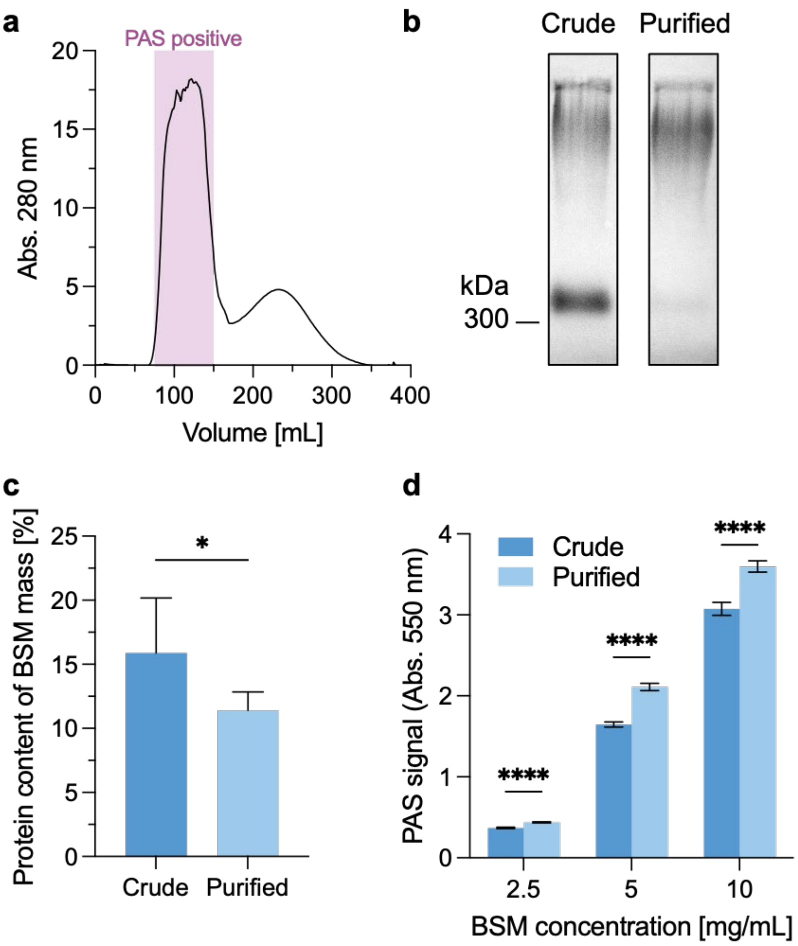


**Figure S1**. Purification of mucins from bovine submaxillary gland (BSM) by size exclusion chromatography and comparison with the crude BSM. (**a**) Size exclusion chromatogram of BSM. Fractions containing mucin were identified by periodic acid/Schiff (PAS) and further enriched. (**b**) Agarose gel electrophoresis of BSM before and after purification. Samples are stained with PAS. Equal masses (30 µg) of samples were loaded. (**c**) Protein content of a 10 mg/mL BSM sample measured with the bicinchoninic acid assay (BCA). Student's t-test is used to compare the means between two groups. (**d**) PAS signal measured as absorbance at 550 nm of the BSM samples before and after purification, at different concentrations of dry mass of BSM. Multiple t-test was used to compare the means of the crude and purified groups. Results are presented as mean values ± standard deviation (SD) of N = 3. $p < 0.05$ (*), $p < 0.01$ (**), $p < 0.001$ (***), $p < 0.0001$ (****)

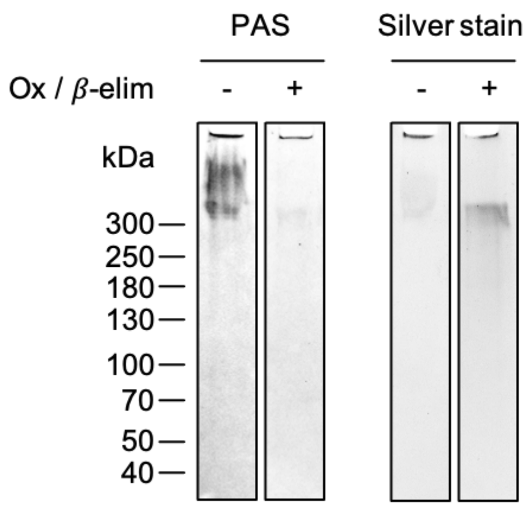


**Figure S2**. Sodium dodecyl sulfate polyacrylamide gel electrophoresis (SDS-PAGE) of purified mucin from bovine submaxillary gland before (-) and after (+) oxidative $\beta$-elimination for glycan removal. Gels are stained with periodic acid-Schiff (PAS) or silver staining to highlight glycans and protein, respectively.

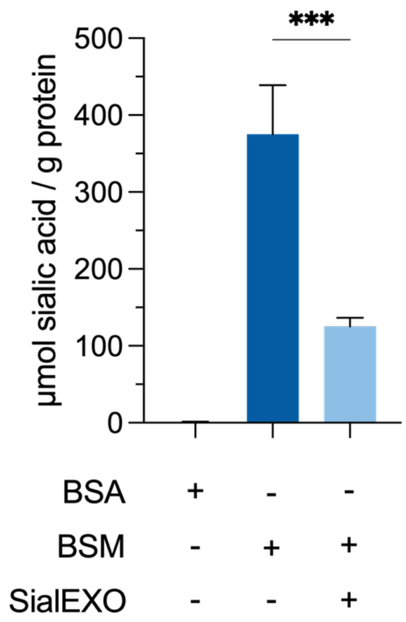


**Figure S3**. Sialic acid quantification after sialidase (SialEXO®) treatment of BSM. Bovine serum albumin (BSA) was used as control of non-glycosylated protein (negative control). Results are reported as mean of N=3 independent measurements ± standard deviation (SD). The significance level was calculated using the Student's t-test. $p < 0.05$ (*), $p < 0.01$ (**), $p < 0.001$ (***), $p < 0.0001$ (****)

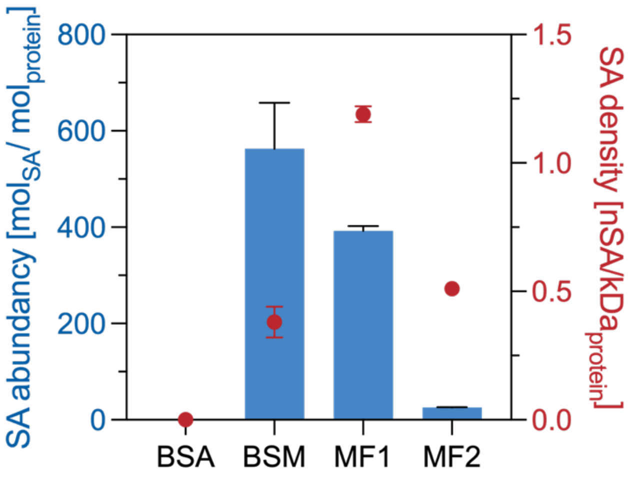


**Figure S4**. Sialic acid abundance (mol of sialic acid per mol of protein) and density (number of sialic acid residues per kDa of protein) in mucin from purified bovine submaxillary mucin (BSM) and in the mucin fragments obtained by StcE (MF1) and proteinase-K (MF2) digestion. Sialic acid abundance was measured using the NANA assay and quantified using a sialic acid five-points calibration curve ($R^2$=0.99). Conversion of mass/volume into molarity concentrations of BSM, MF1, and MF2 was calculated assuming a molecular weight (MW) of 1500, 330, and 50 kDa respectively, according to the apparent MW observed on gel electrophoresis. Bovine serum albumin (BSA) was used as negative control. Results are reported as mean of N=4 independent measurements ± standard deviation (SD).

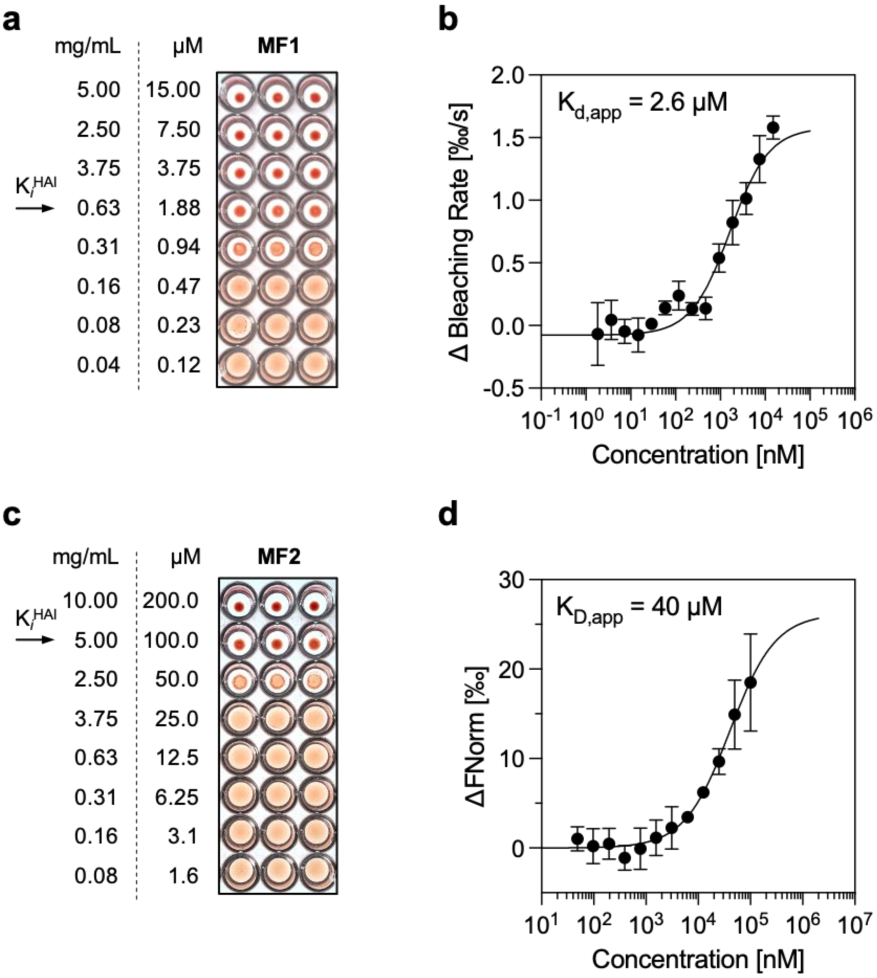


**Figure S5**. Antiviral activity and binding to four hemagglutination units of influenza A/Panama/2007/99 (H3N2) of mucin fragments obtained by digestion with StcE (MF1) and proteinase-K (MF2). Hemagglutination inhibition assay (N=3), and binding curve obtained from microscale thermophoresis analysis of octadecyl rhodamine B chloride (R18) labelled X/31 virus (4 HAU) with MF1 (**a, b**) and MF2 (**c, d)**. Data points were fitted according to the mass-action law function to calculate $K_{D,app}$ values. Results reported as mean of N=4 independent measurements ± standard deviation). Conversion of dry mass/volume into molarity concentrations of MF1 and MF2 was calculated assuming a molecular weight (MW) of 330 and 50 kDa respectively, according to the apparent MW observed on gel electrophoresis (see Figure 5b).

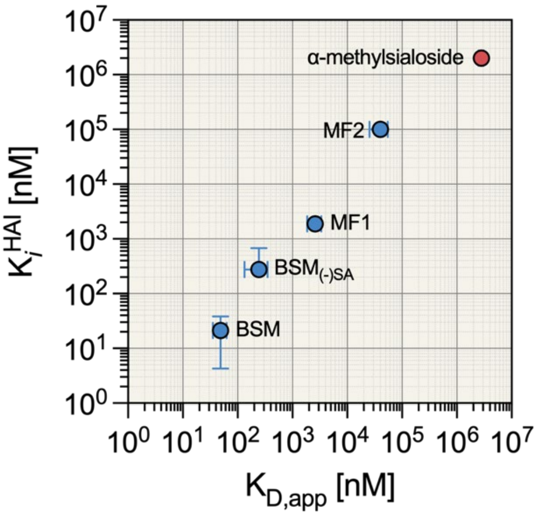


**Figure S6**. Correlation plot between virus inhibition and virus binding. Scatter plot depicting on y-axis the inhibition constant ($K_i^{HAI}$, mean of N=3 ± SD) measured by hemagglutination inhibition assay, and on x-axis the apparent dissociation constant ($K_{D,app}$, mean of N=4 ± SEM) determined from microscale thermophoresis measurements. BSM = mucin from bovine submaxillary gland; $BSM_{(-)SA}$ = BSM after treatment with sialidase to remove sialic acid; MF1 = mucin fragment obtained from StcE digestion; MF2 = mucin fragment obtained from proteinase-k digestion. α-methylsialoside (red dot) is reported from (1) and is used as ligand with minimum affinity and activity in the assay.

### Materials and methods

#### Mucin purification

Mucin from the bovine submaxillary gland (BSM) was obtained from Sigma-Aldrich (Type I-S) as a lyophilized powder. The BSM purification protocol was adapted from a previously published method (2). In short, BSM was dissolved at a concentration of 10 mg/mL in phosphate-buffered saline (DPBS pH 7.4, without calcium and magnesium, Carl Roth), supplemented with sodium chloride at a concentration of 2 M, and stirred overnight at 4 °C. The insoluble fraction was removed by centrifugation (10,000 x g, 4°C, 15 minutes). Subsequently, the mucins were isolated by size exclusion chromatography (SEC) using an Äkta Pure system (Cytiva, Germany) equipped with a HiPrep 26/60 column packed with Sephacryl

400 HR (Cytiva, Germany) and a bed volume of 318.6 mL. DPBS (pH 7.4) supplemented with 2 M sodium chloride was used as equilibration and elution buffer. Approximately 5 mL of the BSM sample was loaded into the column. The loading flow rate was 1 mL/min while the elution flow was set at 0.5 mL/min. Sample elution was recorded by monitoring absorbance at 280 and 214 nm. Fractions of 40 mL were pooled and concentrated by centrifugation filtering (100 kDa MWCO Vivaspin® Turbo 15, Sartorius) and analyzed by periodic acid-Schiff (PAS) reaction to monitor the fractions containing glycosylated material. The concentrated sample was resuspended three times in Milli-Q water to remove the excess salts. The sample was then lyophilized and stored at -80 °C until further use.

**Proteomic analysis**

Crude BSM (#SLCK8402) and three replicates of purified BSM were dissolved in PBS (pH 7.4, without calcium and magnesium) at a concentration of 2 mg/mL. Sample preparation was performed as described by Rulff et al (3). Briefly, 50 µL of the solution was reduced and alkylated by the addition of SDS buffer (2% w/v SDS, 50 mM Tris-HCl pH 8, 0.5 mM EDTA, 75 mM NaCl, 10 mM DTT, 40 mM chloroacetamide final concentration) and incubated at 95 °C for 10 min. After cooling, Benzonase® (10.25 U, Merck) was added and incubated for 15 min at room temperature. The protein sample was purified using a single-pot, solid-phase enhanced sample preparation (SP3 clean-up) (4). Proteins were then incubated for 1 hour with PNGase F (500 U, New England Biolabs) to remove N-glycosylation, followed by overnight digestion with sequence-grade trypsin (Promega) and lysyl endopeptidase (LysC, Wako) (1:50 enzyme:substrate ratio, 37°C). Trifluoroacetic acid was added to a final concentration of 1% (v/v) to stop the digestion. Peptides in the supernatant were desalted using C18 stage tips (5). The eluted and dried peptides were reconstituted in 10 µL of 3% (v/v) acetonitrile (ACN) with 0.1% (v/v) formic acid (FA). 2 µL of peptides were separated on a Vanquish Neo UHPLC system on a reversed-phase column (20 cm fritless silica microcolumns with an inner diameter of 75 µm, in-house packed with ReproSil-Pur C18-AQ 1.9 µm resin (Dr Maisch GmbH)). The gradient length was 98 min at a flow rate of 250 nL/min. The buffer B (90% (v/v) ACN, 0.1% (v/v) FA) concentration was increased from 2% to 60%.

The separated peptides were analyzed on an Orbitrap Exploris 480 mass spectrometer (Thermo Fisher Scientific) in data-dependent acquisition (DDA) mode. At MS1 level, the resolution was 60 k with a normalized AGC target of 300% and a maximum injection time of 10 ms. For MS2 scans, the top 20 ions were selected with a dynamic exclusion time of 30 s. Ions were measured at 15 K resolution with an AGC target of 100% and a maximum injection time of 22 ms.

MaxQuant (V 2.0.3.0) was used for database searching (Uniprot database: bovine proteins (downloaded 2022-09)) (6). Variable modifications were defined as oxidation (M), *N*-terminal acetylation and deamidation (N, Q), and carbamidomethyl (C) were defined as fixed modifications. In addition, "match between runs", label-free quantification, and iBAQ algorithms were used. The mass spectrometry proteomics data have been deposited to the ProteomeXchange Consortium via the PRIDE partner repository with the dataset identifier PXD054384. R (V 4.2.2) was used for statistical analysis. First, proteins were filtered for 'reverse' and 'identified by site only'. Proteins identified as "potential contaminants" were only filtered if they were not from bovine databases. Only proteins quantified in 2 of the 3 purified samples and in the crude BSM sample were considered for further analysis. For statistical analysis, a one-sample *t*-test using the limma-package was performed (7).

**Glycan removal from mucins**

*N*-linked glycans were removed by treatment with peptide N-glycosidase F (PNGase-F) enzyme (New England Biolabs) according to the manufacturer's instructions. Briefly, purified mucin (1 mg/mL) was incubated with PNGase-F (50,000 U/mL) in 5 mM sodium phosphate, pH 7.5, at 37 °C for 24 h. After deglycosylation, the enzyme was inactivated by heating at 75 °C for 10 minutes. The sample was concentrated, and the cleaved *N*-linked glycans were removed by three cycles of centrifugation filtering (50 kDa MWCO Amicon Ultra-4). The structure of the filtered *N*-glycans were further analyzed (see section 2.6.)

Desialylated mucin was prepared by incubation of purified mucin with SialEXO (G1-SM1-020, Genovis) according to the manufacturer's instructions. Shortly, 1 µg of purified mucin was incubated with 1 unit of SialEXO at 37 °C overnight in 20 mM Tris HCl buffer, pH 6.8. The cleaved sialic acid was removed by three cycles of centrifugation filtering (100 kDa MWCO Amicon Ultra-4).

Unspecific deglycosylation was performed following a previously published protocol consisting of oxidation and β-elimination (8). Briefly, a 10 mg/mL of purified BSM sample was solubilized in 0.33 M NaCl at 4°C overnight. Acetic acid was added to 0.1 M, and the pH was adjusted to 4.5 with 1 M NaOH. The oxidation was started by adding ice-cold 200 mM $NaIO_4$ to a final concentration of 100 mM $NaIO_4$. The solution was left to stand in the dark at 4°C overnight. Next, the unreacted periodate was destroyed by adding ½ volume of 400 mM $Na_2S_2O_3$/100 mM NaI/100 mM $NaHCO_3$ (in a 1:1:1 volume ratio). The elimination was started by adding 1 M NaOH to pH 10.5. After standing for 1 h in the cold while maintaining the pH, the solution was dialyzed (100 kDa MWCO) at 4 °C against water. The sample was then lyophilized and stored at -80 °C until further use.

**Enzymatic mucin fragmentation**

For all the digestion reactions, mucin was first denatured, reduced, and alkylated before protease treatment. Purified mucin was dissolved at a concentration of 50 mg/mL in 6 M guanidinium hydrochloride and 5 mM DTT at 37 °C under mild agitation for 1 hour. Then, the sample was alkylated with 20 mM iodoacetamide at room temperature in the dark for 30 minutes. After alkylation, the sample was diluted with 50 mM Tris HCl buffer until the concentration of guanidinium hydrochloride reached 0.3 M. Instead of Tris buffer, 40 mM HCl, pH 2, was used in the reaction with pepsin (Promega, V1959). To increase protease efficiency, specific adjustments were made to the pH and the composition of the Tris buffer. In particular, SmE and StcE digestions were carried out at pH 8, supplemented with 2 mM $ZnCl_2$; ficin (Merck, F4165), bromelain (Merck, B4882), and papain (Merck, 1.07144) digestions were performed at pH 7, supplemented with 2 mM cysteine; trypsin (Merck, T1426) and chymotrypsin (Merck, C4129) digestions were conducted at pH 8, with the addition of 10 mM $CaCl_2$; finally, elastase (Promega, V1891) and proteinase K (Merck, 124568) digestions were carried out at pH 9 and pH 8, respectively. All the proteases were added in a 1:20 protease:BSM ratio, and the digestion was conducted at 37 °C overnight. The reaction was quenched by adding Protease Inhibitor Cocktail (Abcam, ab271306) in a 1:100 volume ratio.

**Isolation and purification of mucin fragments**

Mucin fragments obtained from the proteinase K digestion were enriched and purified by size exclusion chromatography using an Äkta Pure system (Cytiva, Germany) equipped with a Superdex 200 Increase 10/300 GL prepaked column (Cytiva, Germany). DPBS (pH 7.4) was used as equilibration and elution buffer. Approximately 500 µL of the digested sample were loaded into the column. The loading flow rate was 0.75 mL/min while the elution flow was set at 0.25 mL/min. Mucin fragments obtained from StcE digestion were enriched and purified by affinity chromatography using the same FPLC system equipped with a HisTrap HP 1 mL column (Cytiva, Germany). 20 mM Tris buffer (pH 8.0) containing 25 mM imidazole and 500 mM NaCl was used as loading and washing buffer while the same buffer at 500 mM imidazole concentration was used for the elution phase. Approximately 2 mL sample volume was applied at 0.1 mL/min and eluted at 1 mL/min. Sample elution was recorded by monitoring absorbance at 280 and 214 nm.

Fractions of 2 mL were pooled and concentrated by centrifugation filtering (10 kDa MWCO Vivaspin® Turbo 15, Sartorius) and analyzed by periodic acid-Schiff (PAS) reaction to monitor the fractions containing glycosylated material. The concentrated sample was resuspended three times in Milli-Q water to remove the excess salts. The sample was then lyophilized and stored at -80 °C until further use.

SDS-PAGE/PAS was used to analyze the molecular weight of the mucin fragments, and the hydrodynamic radius in PBS was measured by dynamic light scattering (DLS) using a Nanotemper Prometheus Panta device.

Sialic acid was measured using the NANA assay (MAK314, Sigma-Aldrich) where sialic acid is oxidized to formylpyruvic acid which reacts with thiobarbituric acid to form a pink colored product.

**Structural analysis of *O*- and *N*-glycans**

*O*-glycans from intact purified BSM (2 mg/mL in $H_2O$) and digested BSM fractions (2 mg/mL in $H_2O$) were released by reductive $\beta$-elimination (9). 80 µL of each sample were incubated with 0.5 mM NaOH (10 µL) and 5 M $NaBH_4$ (10 µL) at 50 °C for 16 h. Reactions were quenched with acetic acid (15 µL). The solutions were desalted on 400 mg Dowex 50WX8 cation exchange beads (Sigma-Aldrich, USA). Prior to sample loading, the resin was washed with MeOH (3 x 1 mL), and conditioned with 1 M HCl (1 mL), MeOH (1 mL) and $H_2O$ (1 mL). Samples were loaded on the resin and eluted with $H_2O$ (2 x 500 µL). Glycans were extracted using 50 mg Hypercarb SPE cartridges (Thermo Fischer, USA). Cartridges were conditioned with ACN 0.1% (v/v) TFA (1 mL), and 0.1% TFA in $H_2O$. Glycans were loaded on the cartridge and washed with 0.1% TFA in $H_2O$ (3 x 1 mL). Glycans were eluted with 50% ACN 0.1% TFA (4 x 100 µL) and dried in a SpeedVac.

The collected *N*-glycans (see section 2.3.) were lyophilized and subsequently redissolved in 1 M $NaBH_4$ (100 µL). The reaction mixture was incubated at 50 °C for 3 h. The reaction mixture was quenched with acetic acid (25 µL). The solution was desalted on 400 mg Dowex 50WX8 cation exchange beads (Sigma-Aldrich, USA). Prior to sample loading, the resin was washed with MeOH (3 x 1 mL), and conditioned with 1 M HCl (1 mL), MeOH (1 mL) and $H_2O$ (1 mL). The sample was loaded on the resin and eluted with $H_2O$ (2 x 500 µL). Glycans were extracted using 50 mg Hypercarb SPE cartridges (Thermo Fischer, USA). Cartridges were conditioned with ACN 0.1% TFA (1 mL), and 0.1% TFA in $H_2O$. Glycans were loaded on the cartridge and washed with 0.1% TFA in $H_2O$ (3 x 1 mL). Glycans were eluted with 50% ACN 0.1% TFA (4 x 100 µL) and dried in a SpeedVac.

PGC-LC-MS/MS were performed using a SYNAPT G2-Si spectrometer (Waters, U.K.) equipped with an Acquity UPLC system. The glycan alditols were redissolved in 40 µL and 5 µL were injected. Glycan alditols were separated using a 100 × 2.1 mm I.D. PGC column of 5 µm particle size (Hypercarb, Thermo Scientific, U.S.A.) with a linear gradient from 0 to 40% ACN in 10 mM $NH_4HCO_3$ at room temperature at a flow rate of 150 µL/min over 40 min. Analytes were ionized *via* electrospray ionization in negative ion mode with a capillary voltage of 2.8 kV and a source temperature of 150 °C. Fragmentation *via* collision-induced dissociation was performed using collision energy ramps (15 to 120 eV) dependent on the *m/z* of the precursor ion. The MassLynx software (version 2.0.7, Waters) was used for data acquisition and processing. Chromatograms were deconvoluted and integrated with MZmine 3 (10). Glycan structures were manually identified from MS/MS spectra based on diagnostic fragment ions (11, 12) using GlycoWorkBench 2.1 (13).

**Expression and purification of mucinases**

The pET28b-StcE_Δ35-NHis plasmid was supplied by the Carolyn R. Bertozzi lab (Stanford University) with permission from Natalia Strynadka (University of British Columbia), where the plasmid was originally created. The expression and purification were carried out with Junqioa Jia in the Markus Wahl lab (Freie Universität Berlin). The pET28b-StcE_Δ35-NHis plasmid was transformed using electroporation into electrocompetent *E. coli* BL21(DE3) cells in the presence of 50 µg/mL kanamycin. Bacterial cell culture was produced from a single colony and grown at 37°C, shaking, until an $OD_{600}$ of 0.8, when it was induced with 1.5 mL of 0.2 mM IPTG for 1 L of cell culture. The bacterial cell culture was then placed in a shaking incubator at 18 °C and grown overnight. From this point, the bacterial cells are kept continuously on ice or at 4°C conditions. The culture was spun down, supernatant discarded and two complete Mini EDTA-free protease inhibitor cocktail tablets (Roche) were added to 20 mL concentrated bacterial cell pellet and resuspended in 50 mL lysis buffer (20 mM HEPES pH 7.5, 500 mM NaCl) and lysed using Sonopuls Ultrasonic Homogenizer HD (Bandelin) with 65% amplitude for 25 min. The lysed cells were centrifuged at 21,500 x g at 4°C for 60 min. The supernatant was kept for StcE protein purification using the Äkta Pure FPLC system and the HisTrap HP column (GE Healthcare Life Sciences). Washing with 20 column volumes (CV) of lysis buffer plus 20 mM imidazole, the protein was eluted using a linear gradient from 20 mM to 250 mM imidazole over the course of 20 minutes. StcE protein expression was confirmed via SDS-PAGE and StcE-containing fractions were pooled and then concentrated using Amicon Ultra 30 kDa MWCO filters (Millipore Sigma). The protein was further purified using size exclusion chromatography (SEC) FPLC HiLoad Superdex™ S200 16/60 column (GE Healthcare) equilibrated with DPBS buffer. Purified fractions as identified by SDS-PAGE were pooled and again then concentrated using Amicon Ultra 30 kDa MWCO filters (Millipore

Sigma). Purified and concentrated StcE was then flash-frozen in liquid nitrogen and stored at -80 °C.

For protein expression of SmEnhancin (SmE), *E. coli* BL21-CodonPlus (DE3)-RIPL was transformed with pET28a_SmE, which was kindly gifted by Stacy A. Malaker (Yale University), and streaked out on LB agar plates (kanamycin 50 µg/mL, chloramphenicol 34 µg/mL, streptomycin 50 mg/mL). One colony was picked and grown in LB medium (50 µg/mL kanamycin, 34 µg/mL chloramphenicol, 50 mg/mL streptomycin) at 250 rpm, 37 °C, overnight. With this preculture, TB AIM medium (20 mM glucose, 50 µg/mL kanamycin, 34 µg/mL chloramphenicol, 50 mg/mL streptomycin) was inoculated at an $OD_{600}$ of 0.03, grown at 250 rpm, 37 °C, and the exponentially growing culture was aliquoted at an $OD_{600}$ of 2.67 (including 7% DMSO) and snap-frozen in liquid nitrogen. TB AIM medium (30 mM lactose, 11,25 g/L glycerol, 50 µg/mL kanamycin, 17 µg/mL chloramphenicol) was inoculated with an $OD_{600}$ of 0.04 of the thawed bacterial stock and grown at 25 °C, 250 rpm grown for 23h. The culture was harvested and washed two times with lysis buffer (50 mM HEPES, 200 mM NaCl, 2.5 mM $CaCl_2$, 2.5 mM $MgCl_2$, pH 8.0) by centrifugation at 4000 x g for 10 min. The lysis of a bacterial pellet of 100 mL bacterial culture was performed with 2.5% (v/v) DMSO, 2,5% (v/v) n-propanol, 30 mM N-lauroylsarcosine, 1:100 protease inhibitor (Halt™ Protease Inhibitor Cocktail, EDTA-free (100X)) in lysis buffer (total volume 46 mL) at 50% intensity, 50% cycle for 15 minutes (Sonopuls HD 2200 with sonotrode MS73 by Bandelin). The lysate was centrifuged at 12.000 x g for 30 minutes and filtered through a 0.2 µM PES filter and stored on ice for further use.

The protein was purified using an ÅKTA Pure FPLC (Cytiva) with HisTrap column (Cytiva) following a previously published protocol (14).

**Gel electrophoresis and blotting**

Samples were analyzed by sodium dodecyl sulfate polyacrylamide gel electrophoresis (SDS-PAGE). The samples were mixed with 4X loading dye in a ratio of 3:1 and then thermally denatured at 95 °C for 5 minutes. The mixture was briefly centrifuged, and 10 µL were loaded onto a precast polyacrylamide gel (4-20% Mini-PROTEAN® TGX™, Bio-Rad). Additionally, 5 µL of a prestained protein standard solution (ProSieve QuadColor Protein Marker, 4.6-300 kDa, Lonza) was loaded on a separate lane. The gel was run at 100 V in SDS running buffer (25 mM Tris base, 200 mM glycine, 0.1% w/v SDS) for 80 minutes.

After the run, the gel was rinsed with water to remove the excess SDS running buffer, and the gel was subjected to periodic acid-Schiff (PAS) staining for glycan detection as previously reported (15). First, the gel was fixed in 25% (v/v) MeOH and 10% (v/v) acetic acid for 1 hour under gentle shaking. Then, the gel was washed in water for 20 minutes. Oxidation was performed in a 2% periodic acid solution at room temperature for 15 minutes and washed twice with water for 2 minutes. The oxidized gel was stained with Schiff reagent at room temperature, protecting from light, for 40 minutes. The unreacted Schiff reagent was removed by washing the gel with 1% sodium metabisulfite until a clear background was observed. The gel was rinsed with water and imaged with ChemiDoc XRS (BioRad). Proteins detection was achieved using either Der Blaue Jonas solution (GRP, Germany), or silver staining.

The efficiency of enzymatic desialylation and *N*-glycan removal from mucin was verified by lectin blotting. After separation on SDS-PAGE, samples were transferred to nitrocellulose membrane (Carl Roth, GmbH) using a Trans-Blot® Turbo™ Transfer System (Bio-Rad). The membrane was blocked with 3% bovine serum albumin (BSA) in PBST at 4° overnight, washed, and incubated with fluorescently labeled lectins at room temperature for 1 h. Concanavalin A FITC conjugate (Merck) and wheat germ agglutinin Alexa 647 conjugate (ThermoFisher Scientific) were used to specifically stain *N*-linked glycans and sialic acid, respectively. The specificity of the lectin binding was tested using untreated mucin as a positive control, while bovine serum albumin was used as the negative control.

The unpurified and purified mucins were also analyzed using agarose gel electrophoresis and stained by PAS as previously described (16, 17).

**Hemagglutination inhibition assay**

Hemagglutination inhibition assay was conducted using A/Panama/2007/1999 (H3N2) virus as previously described (1). First, the virus was titrated for the hemagglutination assay (HA). Shortly, 50 µL virus concentrate was serially diluted twofold in DPBS using 96-well plates. Then, 50 µL of 1% human red blood cells (German Red Cross, Berlin) were added to each well and incubated at room temperature for 60 minutes. The HA unit (HAU) per 50 µL of virus solution was identified as the last well showing hemagglutination.

The hemagglutination inhibition assay (HAI) was conducted by incubating the virus with human red blood cells to yield agglutination and concentration-dependent inhibition of agglutination (*i.e.*, without and with an agglutination inhibitor). The compounds (*i.e.*, mucin-derived samples) were two-fold serially diluted in DPBS (pH 7.4, without calcium and magnesium), and 4 HAU were added to all wells. The virus and the compounds were incubated at room temperature for 30 minutes to reach equilibrium. After incubation, 50 µL of 1 % human red blood cells were added. The plate was gently tapped and further incubated at room temperature for 1 hour. The lowest inhibitor concentration necessary to achieve complete inhibition of agglutination is defined as the inhibitor constant ($K_i^{HAI}$); this was determined by analysis of the plate after 1 hour of incubation and was expressed as the total dry mass of the compound per volume (µg/mL).

**Microscale thermophoresis**

Binding measurements of the influenza virus were carried out by microscale thermophoresis (MST) using a Monolith NT.115 (Nanotemper) from the Heberle lab (FU Berlin). The envelope of a 1 mg/mL (expressed as protein content) suspension of A/X31/1 (A/Aichi/1968 (H3N2) reassorted with A/Puerto Rico/8/1934 (H1/N1)) virus was labeled with 20 µM octadecyl rhodamine B (R18, Invitrogen), under gentle shaking on ice for 30 min. Unbound R18 was removed using desalting columns (Zeba™7K MWCO, Thermo Scientific). The collected virus was filtered using a 0.45 µm filter to remove virus aggregates. As a quality control, the amount of virus and its binding ability were tested by performing a hemagglutination assay. All MST measurements were carried out at 22 °C in premium capillaries using default settings (initial fluorescence = 5s, thermophoresis = 30s, and recovery = 5s). The MST power was set at 20% and the LED power (green LED) at 80%. For affinity measurement, the ligands were serially diluted 1:2 in PBS and mixed with R18-labeled virus (4 HAU, C final ~0.1 nM virus particles). The obtained data for the mucin building block was analyzed with a gating strategy 1.5 s after the start of the thermophoresis, and fitted as previously shown (1). The data obtained for intact mucin was analyzed based on the initial fluorescence and fitted according to the bleaching rate, as previously shown (18).

### References


1. S. Bhatia, *et al.*, Linear polysialoside outperforms dendritic analogs for inhibition of influenza virus infection in vitro and in vivo. *Biomaterials* **138**, 22–34 (2017).
2. V. J. Schömig, *et al.*, An optimized purification process for porcine gastric mucin with preservation of its native functional properties. *RSC Adv.* **6**, 44932–44943 (2016).
3. H. Rulff, *et al.*, Comprehensive Characterization of the Viscoelastic Properties of Bovine Submaxillary Mucin (BSM) Hydrogels and the Effect of Additives. *Biomacromolecules* (2024). https://doi.org/10.1021/acs.biomac.4c00153.
4. C. S. Hughes, *et al.*, Ultrasensitive proteome analysis using paramagnetic bead technology. *Mol. Syst. Biol.* **10**, 757 (2014).
5. J. Rappsilber, Y. Ishihama, M. Mann, Stop and Go Extraction Tips for Matrix-Assisted Laser Desorption/Ionization, Nanoelectrospray, and LC/MS Sample Pretreatment in Proteomics. *Anal. Chem.* **75**, 663–670 (2003).
6. S. Tyanova, T. Temu, J. Cox, The MaxQuant computational platform for mass spectrometry-based shotgun proteomics. *Nat. Protoc.* **11**, 2301–2319 (2016).
7. M. E. Ritchie, *et al.*, limma powers differential expression analyses for RNA-sequencing and microarray studies. *Nucleic Acids Res.* **43**, e47–e47 (2015).
8. T. A. Gerken, R. Gupta, N. Jentoft, A novel approach for chemically deglycosylating O-linked glycoproteins. The deglycosylation of submaxillary and respiratory mucins. *Biochemistry* **31**, 639–648 (1992).
9. P. H. Jensen, N. G. Karlsson, D. Kolarich, N. H. Packer, Structural analysis of N- and O-glycans released from glycoproteins. *Nat. Protoc.* **7**, 1299–1310 (2012).
10. R. Schmid, *et al.*, Integrative analysis of multimodal mass spectrometry data in MZmine 3. *Nat. Biotechnol.* **41**, 447–449 (2023).
11. A. V Everest-Dass, J. L. Abrahams, D. Kolarich, N. H. Packer, M. P. Campbell,

Structural Feature Ions for Distinguishing N- and O-Linked Glycan Isomers by LC-ESI-IT MS/MS. *J. Am. Soc. Mass Spectrom.* **24**, 895–906 (2013).

12. M. Grabarics, *et al.*, Mass Spectrometry-Based Techniques to Elucidate the Sugar Code. *Chem. Rev.* **122**, 7840–7908 (2022).

13. A. Ceroni, *et al.*, GlycoWorkbench: A Tool for the Computer-Assisted Annotation of Mass Spectra of Glycans. *J. Proteome Res.* **7**, 1650–1659 (2008).

14. M. A. Rosenfeld, *et al.*, Glycoproteomic landscape and structural dynamics of TIM family immune checkpoints enabled by mucinase SME and molecular dynamics simulations. *Biophys. J.* **123**, 14a (2024).

15. K. Jiang, *et al.*, Modulating the Bioactivity of Mucin Hydrogels with Crosslinking Architecture. *Adv. Funct. Mater.* **31**, 2008428 (2021).

16. K. A. Ramsey, Z. L. Rushton, C. Ehre, Mucin Agarose Gel Electrophoresis: Western Blotting for High-molecular-weight Glycoproteins. *J. Vis. Exp.* (2016). https://doi.org/10.3791/54153.

17. D. J. Thornton, J. K. Sheehan, I. Carlstedt, “Identification of Glycoproteins on Nitrocellulose Membranes and Gels BT - Basic Protein and Peptide Protocols” in J. M. Walker, Ed. (Humana Press, 1994), pp. 119–128.

18. J. A. C. Stein, A. Ianeselli, D. Braun, Kinetic Microscale Thermophoresis for Simultaneous Measurement of Binding Affinity and Kinetics. *Angew. Chemie Int. Ed.* **60**, 13988–13995 (2021).
